# Embryonic signals mediate extracellular vesicle biogenesis and trafficking at the embryo–maternal interface

**DOI:** 10.1101/2022.11.16.516738

**Authors:** Maria M. Guzewska, Kamil Myszczynski, Yael Heifetz, Monika M. Kaczmarek

**Affiliations:** Department of Hormonal Action Mechanisms, Polish Academy of Sciences, Olsztyn, Poland; Molecular Biology Laboratory, Institute of Animal Reproduction and Food Research, Polish Academy of Sciences, Olsztyn, Poland; The Department of Entomology, The Hebrew University of Jerusalem, Jerusalem, Israel

**Keywords:** extracellular vesicles, pregnancy, implantation, embryo, endometrium, embryonic signals, estradiol, prostaglandins, microRNA

## Abstract

Extracellular vesicles (EVs) are membrane-coated nanoparticles secreted by almost all cell types in living organisms. EVs, as paracrine mediators, are involved in intercellular communication, immune response, and several reproductive events, including the maintenance of pregnancy. Using a domestic animal model (*Sus scrofa*) with an epitheliochorial, superficial type of placentation, we focused on EV biogenesis pathway at the embryo–maternal interface, when the embryonic signaling occurs for maternal recognition and the maintenance of pregnancy. Transmission electron microscopy was used during early pregnancy to visualize different populations of EVs and apocrine and/or merocrine pathways of secretion. Immunofluorescent staining localized proteins responsible for EV biogenesis and cell polarization at the embryo–maternal interface. The expression profiles of genes involved in biogenesis and the secretion of EVs pointed at the possible modulation of endometrial expression by embryonic signals. Further in vitro studies showed that factors of embryonic origin can regulate the expression of the ESCRT-II complex and EV trafficking in luminal epithelial cells. Moreover, miRNA-mediated rapid negative regulation of gene expression was abolished by delivered embryonic signals. Our findings demonstrated that embryonic signals are potent modulators of EV-mediated secretory activity of the endometrium during the critical stages of early pregnancy.

## 1 INTRODUCTION

The establishment of cell-to-cell communication between the endometrium and developing embryo is one of the most important steps in successful pregnancy [1]. This close interaction between the uterine luminal epithelium and trophoblast cells requires common among eutherian mammals targeted apposition and adhesion to initiate either invasive or noninvasive placentation [2]. During this period most pregnancy losses occur in most of mammals, including humans [3]. A crucial point in ensuring that pregnancy can be carried to full term is the maternal recognition of pregnancy—when the embryo signals its presence in the uterine lumen. This stage of pregnancy is manifested by a species-specific signals production and their interaction with receptors. Although mechanism of the maternal recognition of pregnancy differs between species it is equally needed to support corpus luteum maintenance for continued production of progesterone required for secretory activity of the endometrium essential for embryonic development, implantation and placentation [4]. Among hundreds of molecules known to be involved in the complexity of the embryo–maternal interaction, several conserved pathways have been identified. [2].

Pregnancy establishment in the pig is mediated mainly by 17β-estradiol (E_2_), secreted by conceptuses in biphasic manner on days 11–12 and then 15–30 of pregnancy [5]. Estradiol can regulate the differentiation, proliferation, and secretory activity of epithelial cells induced by the interactions with specific receptors, followed by alterations in gene expression important in the initiation of implantation and the formation of the epitheliochorial placenta [6]. The additional abundant production of other important mediators, such as prostaglandin E_2_ (PGE_2_) occurs. PGE_2_ is synthesized and secreted by conceptuses and endometrium, stimulating the endometrial secretion of PGE_2_ and the expression of its receptors—PTGER2 and PTGER4 [7, 8]. Thus far, numerous studies have been conducted to investigate the effects of embryonic signaling (E_2_ and PGE_2_) on the gene expression profile in the porcine endometrium during the estrous cycle and pregnancy [9]; however, only a few have looked at non-transcriptional and non-translational adjustments governed by extracellular vesicles (EVs).

A growing body of evidence emphasizes the importance of EVs as mediators of embryo–endometrial cross talk during early pregnancy [10–12]. EVs were observed for the first time at the porcine embryo– maternal interface as numerous, small vesicles between the microvilli of the luminal epithelium and trophoblast cells [13]. It is now clear that EVs carry lipids, proteins, enzymes, as well as hormones and nucleic acids (e.g., microRNAs [miRNA]). Due to their small size, EVs can cross the intercellular physical barrier and play a significant role in intercellular communication [14], which is crucial at the embryo– maternal interface. EV biogenesis is a dynamic process, governed by various checkpoints. EVs released from cells are a heterogeneous population of nanoparticles, which, according to the guidelines of MISEV 2018, can be distinguished based on physical aspects (small or medium/large vesicles) or biochemical composition (cargo or presence of positive markers) [15]. Small EVs known as intraluminal vesicles (ILVs), are formed by inward membranes budding into the late endosomes and creating multivesicular bodies (MVBs). The endosomal sorting complex required for transport (ESCRT) was the first discovered mechanism involved in the formation of MVBs, catalyzing one of the most unusual membrane remodeling processes in the cell [16]. The ESCRT complex is composed of four different complexes—ESCRT-0, -I, -II, and -III—supported by associated proteins like the programmed cell death 6 interacting protein (Alix) and ATPase (vacuolar protein sorting 4, VPS4) [17]. Furthermore, ILVs can be formed during ESCRT-independent pathway involving ceramide generation [18, 19]. Medium/large EVs bud directly out of the plasma membrane. In addition to the ESCRT complex, several Ras-related proteins, governing secure trafficking in the endocytic and secretory pathways (e.g., Rab27A, Rab7B, and Rab11A/B) and intercellular cell-to-cell communication, have been implicated in the regulation of small and medium/large EVs release, as well as cell polarization [20, 21]. Despite the constant broadening of knowledge regarding EVs, the mechanism responsible for the selection of a particular biogenesis pathway remains unknown.

It has been proposed that EVs secreted from the luminal epithelium might influence the conceptus to migrate to the implantation site and attach to the luminal epithelium [11]. Trophoblast cell polarization and the receptivity of the endometrial luminal epithelial cells can be altered by non-transcriptional, non-translational mechanisms including the exchange of signaling molecules via EV-dependent pathways [22, 23]. To further understand the spatiotemporal dynamics of cell-to-cell communication during early pregnancy, we tracked EV biogenesis components at the embryo– maternal interface in pigs. Here, we investigated the impact of embryonic signaling on the ability of the uterine endometrium to generate and secrete EVs. Our results showed dynamic patterns of ESCRT complex and Rab gene expression at the embryo–maternal interface, which corresponded with the E_2_ secretion by the conceptuses. Treatment of the uterine luminal epithelium with E_2_ altered ESCRT complex gene expression in the endometrium, which may affect EV biogenesis during embryo implantation. Finally, fast inhibition of gene expression [24] was evident after delivery of miRNA to luminal epithelial cells, which was, however, abrogated by embryonic signals. Our data highlight the way that signals present in the uterine environment (e.g., non-coding RNAs and hormones released by the embryo and/or uterus) can be involved in the EVs biogenesis regulation, formation and secretion at the embryo–maternal interface.

## 2 MATERIALS AND METHODS

### 2.1 Animals

Crossbred or cyclic gilts (Hampshire × Duroc) of similar age (7-8 months) and weight (140 ± 150 kg) were used for all experiments. After their second natural estrus, gilts were randomly assigned to a group of cyclic or pregnant animals; the latter were artificially inseminated and again 24 h after onset of estrus (first day of pregnancy [DP]). For transmission electron microscopy we used tissues collected from animals on day 12 (n = 3), 16 (n = 3), and 20 (n = 3) of pregnancy after flushing uterine horns with 20 ml 0.01 M PBS (Phosphate-buffered saline; pH 7.4) to remove conceptuses. For mRNA isolation and immunofluorescent staining, tissues (embryos/endometrium) were collected on 11-12 (n = 13/5), 15-16 (n =8/9), and 17-19 (n = 5/5) days of the estrous cycle/pregnancy. The day of pregnancy was confirmed by size and morphology of conceptuses: days 11-12 (from tubular to filamentous > 100 mm long), 15-16 (elongated), 17-19 (presence of trophoblast tissue and well-developed embryos) [12]. Obtained conceptuses were immediately fixed in *10%* buffered formalin (Neutral Phosphate Buffered Formalin Fixatives, Leica, Germany) or frozen in liquid nitrogen and stored at −80 °C. The same procedure was performed for endometrial tissues. Additionally, several sections (2-3 cm long) from not flushed uterine horns were fixed in fresh 10% buffered formalin. Uteri collected from gilts on days 11-12 of the estrous cycle (DC) served for isolation of endometrial luminal epithelial cells.

### 2.2 Transmission electron microscopy

Fragments of uterine tissue and conceptuses were fixed in fresh solution containing 2.5% glutaraldehyde in 0.1 M cacodylate buffer (pH 7.4) for 12 h at 4 °C. The tissues were then rinsed in 0.1 M cacodylate buffer (pH 7.4), post fixed and stained with 1%osmium tetroxide, 1.5%potassium ferricyanide in 0.1 M cacodylate buffer for additional 2 h at room temperature followed by embedding in epoxy resin. Ultrathin sections (80 nm) of embedded tissues were sectioned with a diamond knife on a Ultracut E microtome (Reichert Jung, Germany) and collected onto 200 Mesh, thin bar copper grids. Sections on grids were sequentially stained with Uranyl acetate for 10 min and Lead citrate for 10 min. Randomly selected sections (10 per animal/tissue type) of endometrial (luminal epithelium/glandular epithelium) or embryonic tissue were captured with a Tecnai 12 transmission electron microscope (FEI), operating at an acceleration 100 kV and equipped with a CCD camera MegaView II and Analysis version 3.0 software (SoftImaging System GmbH, Germany). Finally, 296 images (12 DP = 133; 16 DP = 94; 20 DP = 69) were subjected to further image analysis and interpretation.

### 2.3 Immunofluorescent tissue staining and confocal imaging

After fixation in 10% buffered formalin for 24-48 h and embedding into paraffin, tissues were cut into 4 μm thick sections and mounted on chromogelatin-precoated slides (Leica, Germany). After paraffin removal, tissue was dehydrated in ethyl alcohol grades (100-70%). Non-specific antigens were blocked for 60 min using a blocking buffer (Sea Block Blocking Buffer; ThermoFisher Scientific, USA). Then, slices were incubated for 20 min with 0.3%Sudan Black (Sigma-Aldrich, USA) in the dark. After incubation, sections were washed with a TBS buffer (50 mM TrisHCl, pH 7.4; 150 mM NaCl). Incubation with primary antibodies (Supplementary Table 1) was performed overnight at 4°C. Afterwards, slides were washed in TBS buffer and incubated for 1 h with secondary antibodies (Supplementary Table 1). Finally, sections were mounted in Ultra Cruz Mounting Medium (Santa Cruz Biotechnology, USA) containing 4′,6-Diamidino-2-Phenylindole (DAPI). Negative controls were performed without primary antibodies (Supplementary Fig. 1A, B). From randomly chosen sections, 10-15 images were captured by Laser Scanning Microscope (LSM800; Carl-Zeiss, Germany) equipped with AiryScan super-resolution module using 40×/1.2 NA c-apochromat objective and further analyzed in ZEN Blue Pro 2.6 software (Carl-Zeiss, Germany).

### 2.4 Isolation of endometrial luminal epithelial cells

Porcine endometrial luminal epithelial cells were isolated according to the method described by Blitek & Ziecik [25], with modifications. Briefly, uterine horns collected from gilts on day 11-12 of the estrus cycle (n = 4 - 7) were washed three times in sterile 0.01 M PBS (pH 7.4). Endometrium was separated from the myometrium and digested with 0.2% Dispase II (Sigma-Aldrich, USA) in HBSS (Ca-, Mg- and phenol red-free; pH 7.4; Sigma-Aldrich, USA) at room temperature for 45 min, using a magnetic stirrer. Released luminal epithelial cells were centrifuged at 200 × *g* for 8 min at 4 °C, then washed with Medium 199 (M199; Sigma-Aldrich, USA) containing 10% Newborn Calf Serum (NCS, Sigma-Aldrich, USA) and 100 IU/ml penicillin and 100 g/ml streptomycin (Sigma-Aldrich, USA). Blood cells were removed using the Red Blood Cell Lysing Buffer (Sigma-Aldrich, USA). After final centrifugation, the obtained pellet was suspended M199 containing 10% NCS and antibiotics and pleated onto 6-well plates (Corning, USA) or onto Cell Imaging Coverglasses (Eppendorf, Germany) and then cultured in M199 containing 10% NCS until reaching 90 - 100%confluence, before the beginning of a treatment.

### 2.5 In vitro cell culture and treatments

#### Hormone treatment

Since PGE_2_ and E_2_ act together as embryonic signals during the first days of pregnancy [7], its action was tested on primary luminal epithelial cells collected from gilts on 11-12 day of the estrous cycle, not exposed to the embryonic signals. In a preliminary study, we used two different doses of E_2_ (10 nM, 100 nM; Sigma-Aldrich, USA) and PGE_2_ (1 uM, 100 nM; Cayman Chemicals, USA). Based on this study, only one dose - 100 nM for both factors was used in subsequent experiments (Supplementary Fig. 2A). In addition, methyl-piperidinopyrazole (MPP, Sigma-Aldrich, USA), the most selective estrogen receptor alpha (ERα or ESR1) antagonist [26] was used, as porcine luminal epithelial cells are devoid of beta receptor (ERβ or ESR2; Supplementary Fig. 2B). For PGE_2_, AH6809 (Sigma-Aldrich, USA), a PTGER2 antagonist was selected as a sufficient inhibitor of the main receptor mediating its action on day 11-12 of pregnancy in pigs [7]. To investigate the effect of PGE_2_ on EV biogenesis pathway, luminal epithelial cells were cultured in serum-free M199 (control) or pretreated with AH6809 (10 μM) for 10 min [7]. Subsequently, cells were incubated with PGE_2_ (100 nM in serum-free M199) alone or in the presence of AH6809. To test the effect of E_2_, cells were cultured in serum-free M199 (control) or pretreated with MPP (1 μM) for 1 h [27]. Subsequently, cells were incubated in culture medium supplemented with E_2_ (100 nM in serum-free M199) alone or in the presence of MPP. In addition, simultaneous treatment with both embryonic signals was tested (100 nM E_2_ and 100 nM PGE_2_).

#### miRNA and hormone treatment

Considering the presence [12] and important role of miRNAs in pregnancy establishment and trophoblast differentiation, proliferation and migration [28] we tested miRNA and hormone interplay in regulation of EVs biogenesis pathway. Chosen mimics, miR-125b-5p (MIMAT0000423; 50nM [28]) or control mimic (50nM; Life Technologies, USA, Supplementary Table 2), were delivered to primary luminal epithelial cells collected from gilts on 11-12 day of the estrous cycle, using Lipofectamine RNAiMAX (Life Technologies, USA). Cells were incubated for 12 h with mimics before or after embryonic signals (100 nM E_2_ and 100 nM PGE_2_; incubation for 24 h).

All in vitro treatments were performed at 37 °C in a humidified atmosphere of 95 % air to 5 %CO_2_. Afterwards, culture medium was removed and luminal epithelial cells were washed in PBS, lysed in TRI Reagent (Invitrogen, USA) and subjected to total RNA isolation.

### 2.6 Immunofluorescent cell staining and confocal imaging

To investigate the effect of embryonic signals and miRNAs on the EV biogenesis pathway proteins, immunofluorescent staining was performed. Luminal epithelial cells were cultured in Cell Imaging Coverglasses (Eppendorf, Germany). Cells subjected to following treatments (n = 3-5) as described above were fixed in 4%buffered formalin, permeabilized in 0.25% Triton X-100 and blocked in Sea Block Blocking Buffer (1 h; ThermoFisher Scientific, USA). Overnight incubation with primary antibodies was followed by the incubation (1 h) with Cy3 or Alexa Fluor 488 conjugated secondary antibodies (Supplementary Table 1). At the end of incubation, cells were treated with DAPI solution (0.1 mg/mL; Sigma-Aldrich, USA) to visualize nucleus and with phalloidin conjugated with Alexa Fluor 488 (200 units/mL; ThermoFisher Scientific, USA), in order to visualize cytoskeleton in control staining. Negative controls were performed without primary antibodies (Supplementary Fig. 1C).

Samples were visualized using confocal Laser Scanning Microscope (LSM800; Carl-Zeiss, Germany) equipped with AiryScan super-resolution module using 40×/1.2 NA c-apochromat objective. Ten images of luminal epithelial colonies, showing of at least 10 well-visible cells were captured for each animal. Regions of interests (ROIs) were designated by the eye-based evaluation of the quantity and quality of the luminal epithelial cells in monolayer, as well as the localization of both protein markers (VPS36 and CD63) at the edges of cells. Next, VPS36 and CD63 fluorescence intensity as well as theirs colocalization coefficiency (CC) were analyzed using ZEN Blue Pro 2.6 software (Carl-Zeiss, Germany). CC indicates the relative number of colocalized pixels in relation to the total number of pixels above the image background-noise threshold value, which was constant during analysis.

### 2.7 Total RNA isolation and real-time PCR

Total RNA was isolated using RNeasy Mini Kit (Qiagen, Germany) according to the standard manufacturer protocol and stored in −80 °C for further real-time PCR (qRT-PCR) analysis. Gene expression was assessed using TaqMan Gene Expression Assays (Supplementary Table 3), TaqMan RNA–to Ct1-Step Kit (ThermoFisher Scientific, USA) and Applied Biosystems Fast Real-time PCR system 7900HT according to the manufacturer’s protocol (ThermoFisher Scientific, USA). In each qRT-PCR reaction (performed in duplicates), 15 ng of total RNA was used as a template. Negative controls without a template were performed in each run. The expression values were calculated including the efficiency of the reactions and normalized to reference genes selected among GAPDH, HPRT1, ACTB-1 using NormFinder [29].

### 2.8 In silico screening of binding motifs

To identify estrogen response elements (ERE) and putative prostaglandin-response elements (PGREs) within promoter regions of chosen genes, the 3000 bp sequence upstream and 200 bp downstream from the transcription starting site (TSS) was extracted. The specified region was scanned for matches (allowing two possible mismatches) using FIMO within MEME Suite 5.1.0 [30].

### 2.9 Statistical analysis

Statistical analysis was carried out using GraphPad Prism 9.10 (GraphPad Software Inc., USA). Effects were considered significant at p < 0.05. Logarithmic transformation of the data was performed for samples without a normal distribution (Shapiro-Wilk test). Statistical tests were selected based on the experimental set up and included two-way ANOVA followed by Sidak’s post hoc test (reproductive status [rep] = pregnancy/estrous cycle and day [day] as main factors), one-way ANOVA followed Tukey’s post hoc test or Kruskal-Wallis and Dunn’s post hoc test, and paired T-test or Wilcoxon matched-pairs signed rank test. Unpaired T-test was used for immunofluorescence intensity evaluation and Pearson correlation coefficient (PCC) was used for quantitative analysis of VPS36 and CD63 immunocolocalization [31]. Sample sizes and other statistical details are indicated in the figures/figure legends.

## 3 RESULTS

### 3.1 Dynamic biogenesis and exchange of EVs at the embryo–maternal interface is evident during early pregnancy

To examine cell-to-cell communication at the embryo–maternal interface, we used transmission electron microscopy to screen EV biogenesis and trafficking in endometrial and trophoblast tissues collected on the following days of pregnancy (DP) during the early stages of embryo implantation: day 12 (12 DP; maternal recognition of pregnancy, embryo apposition), day 16 (16 DP; initial stages of embryo attachment), and day 20 (20 DP; firm embryo attachment; Figs. 1–3, Supplementary Fig. 3).

**Figure 1.**
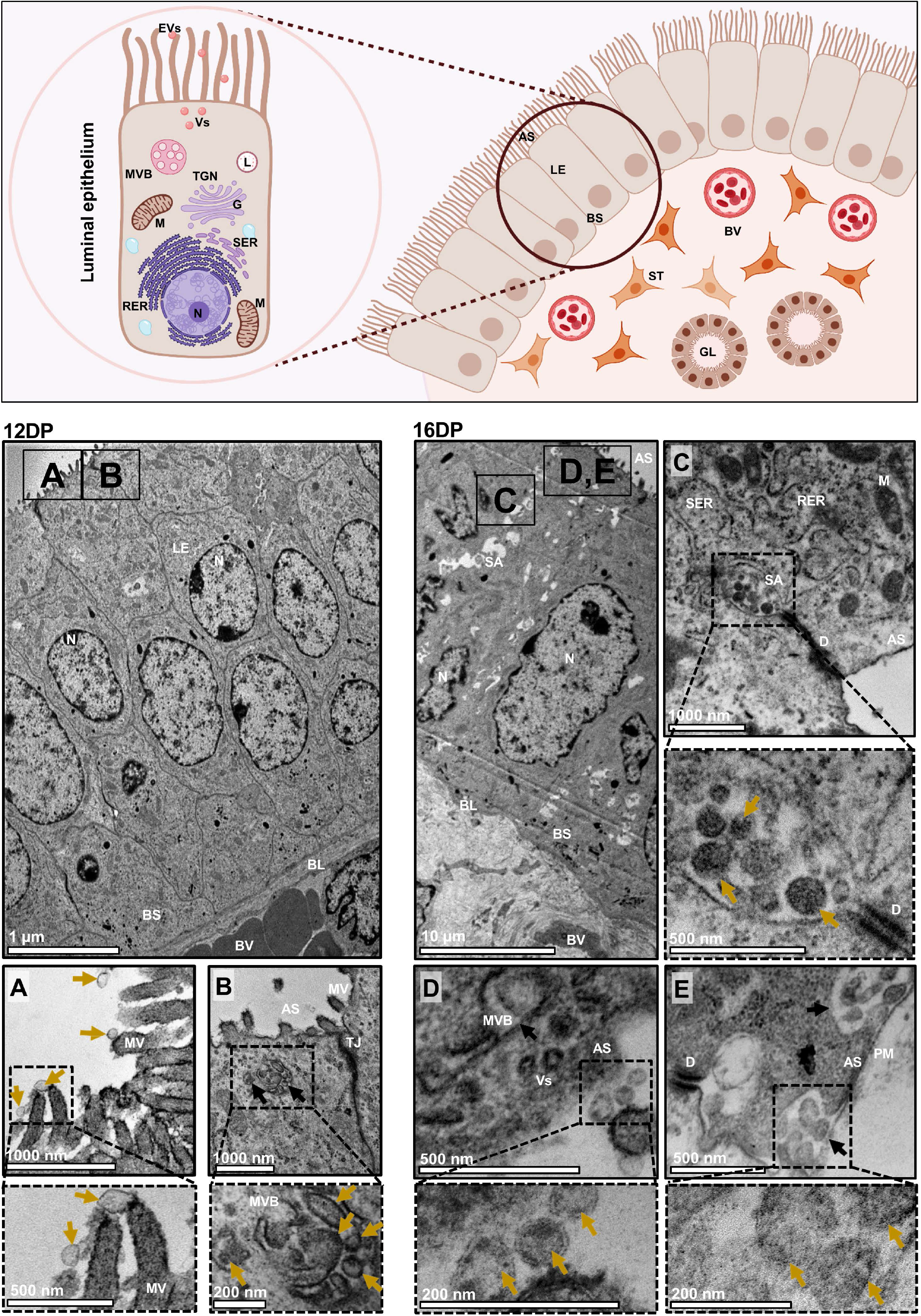
Dynamic biogenesis and release of EVs in the endometrial luminal epithelium during early pregnancy. *Upper panel:* Schematic representation of the superficial endometrium. Uterine luminal epithelium is composed of the population of polarized cells. Apical surface of luminal epithelial cell is facing the uterine lumen. The basal lamina is located close to the complex stromal compartment filled with different cells and structures, including stromal cells, blood vessels, and glands. *Left panel:* Cross sections of luminal epithelium on day 12 of pregnancy showing tall columnar epithelial cells with well-developed microvilli on the apical surface. **(A)** A large population of EVs (yellow arrows) docking on the microvilli was observed. **(B)** MVBs (black arrows) were located at the apical site of the cell. MVBs contained inside a population of heterogeneously sized EVs (higher magnification). *Right panel:* Cross sections of luminal epithelium on day 16 of pregnancy showing tall columnar epithelium with well-developed microvilli. **(C)** Characteristic secretory areas, containing vesicles were located close to the desmosome at the apical site of the luminal epithelial cells. **(D)** EVs (yellow arrows) were still docking on the microvilli. **(E)** MVBs (black arrows) were found at the apical site of the cell, fused with plasma membrane. MVBs contained inside a population of EVs ready to be released into the extracellular space (higher magnification). AS – apical surface, BL – basal lamina, BS – basal surface, BV – blood vessel, D – desmosome, DP – day of pregnancy, G – Golgi apparatus, L – lysosome, LE – luminal epithelium, M – mitochondria, MV – microvilli, MVBs – multivesicular bodies, N – nucleus, PM – plasma membrane, RER – rough endoplasmic reticulum, SA – secretory areas, SER – smooth endoplasmic reticulum, ST – stromal cell, TJ – tight junction, TGN – Trans Golgi network, Vs – vesicles.

**Figure 2.**
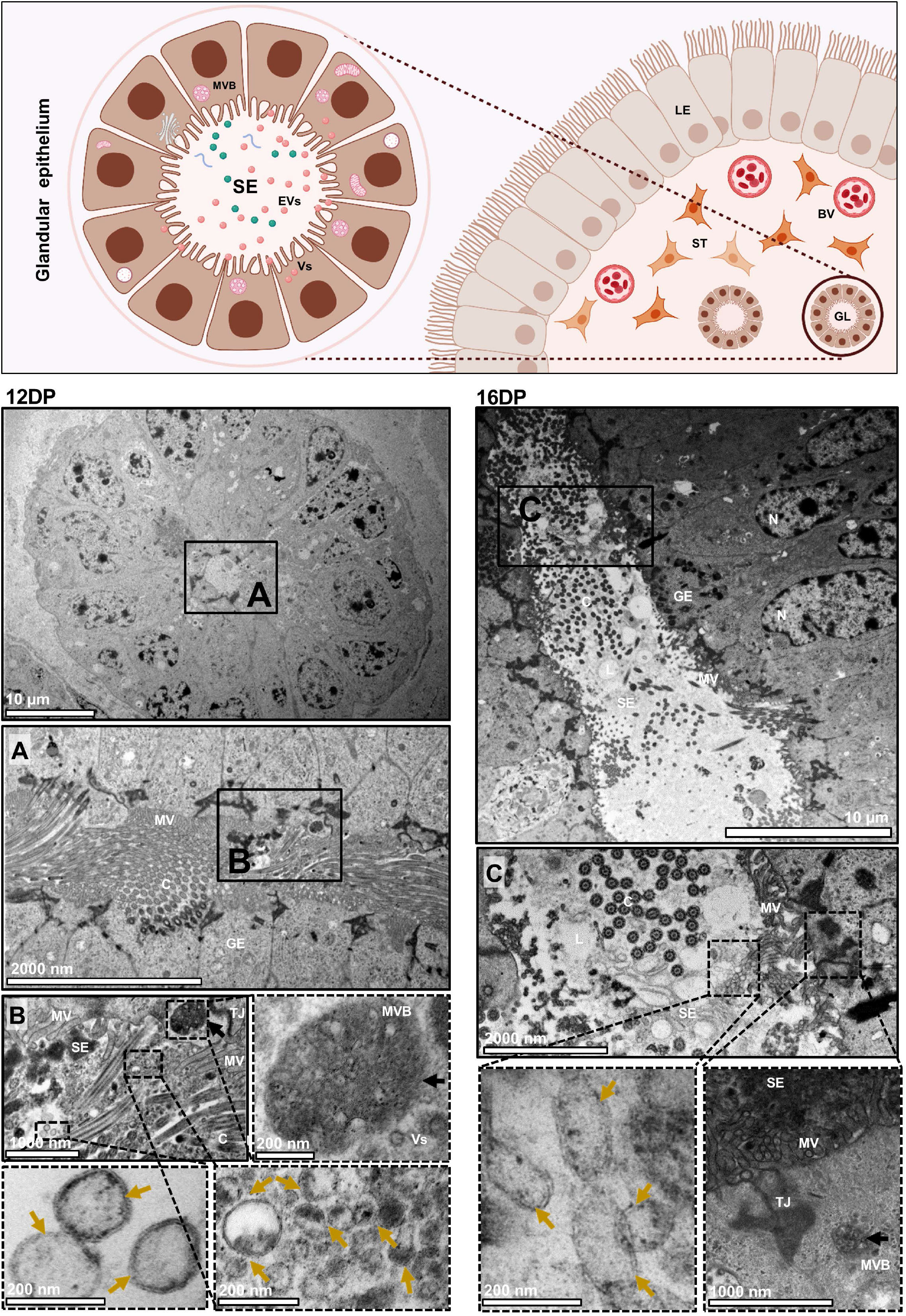
Dynamic change in EV-mediated secretory activity of the glandular epithelium during early pregnancy. *Upper panel:* Schematic representation of the uterine glands. Uterine glands are located in the stromal compartment. Glands are composed of columnar epithelial cells covered with microvilli and cilia. Secretome is present in the glandular lumen and contain population of EVs. MVBs are located at the apical site of glandular epithelial cells, releasing EVs into the lumen. *Left panel:* Cross section of closed uterine gland on day 12 of pregnancy. **(A)** Glands had small diameter of the lumen and were composed of cells covered with cilia and microvilli (higher magnification). **(B)** MVBs (black arrows) were located under the plasma membrane and contained large amount of vesicles (higher magnification). Lumen of the gland was filled with heterogeneous in size population of EVs (yellow arrows) and numerous cilia and microvilli. *Right panel:* Cross section of wide open uterine gland on day 16 of pregnancy. **(C)** Large population of glandular cells with the cilia and the microvilli were observed. Glands were filled with secretome, containing heterogeneous population of EVs (yellow arrows), cached between the microvilli. The apical site of the glandular epithelial cells showed MVBs (black arrow) filled with vesicles located under the plasma membrane (higher magnification). BV – blood vessel, C – cilia, DP – day of pregnancy, EVs – extracellular vesicles, GE – glandular epithelium, L – lumen, MV – microvilli, MVBs – multivesicular bodies, N – nucleus, SE – secretome, ST – stromal cells, TJ – tight junction, Vs – vesicles.

**Figure 3.**
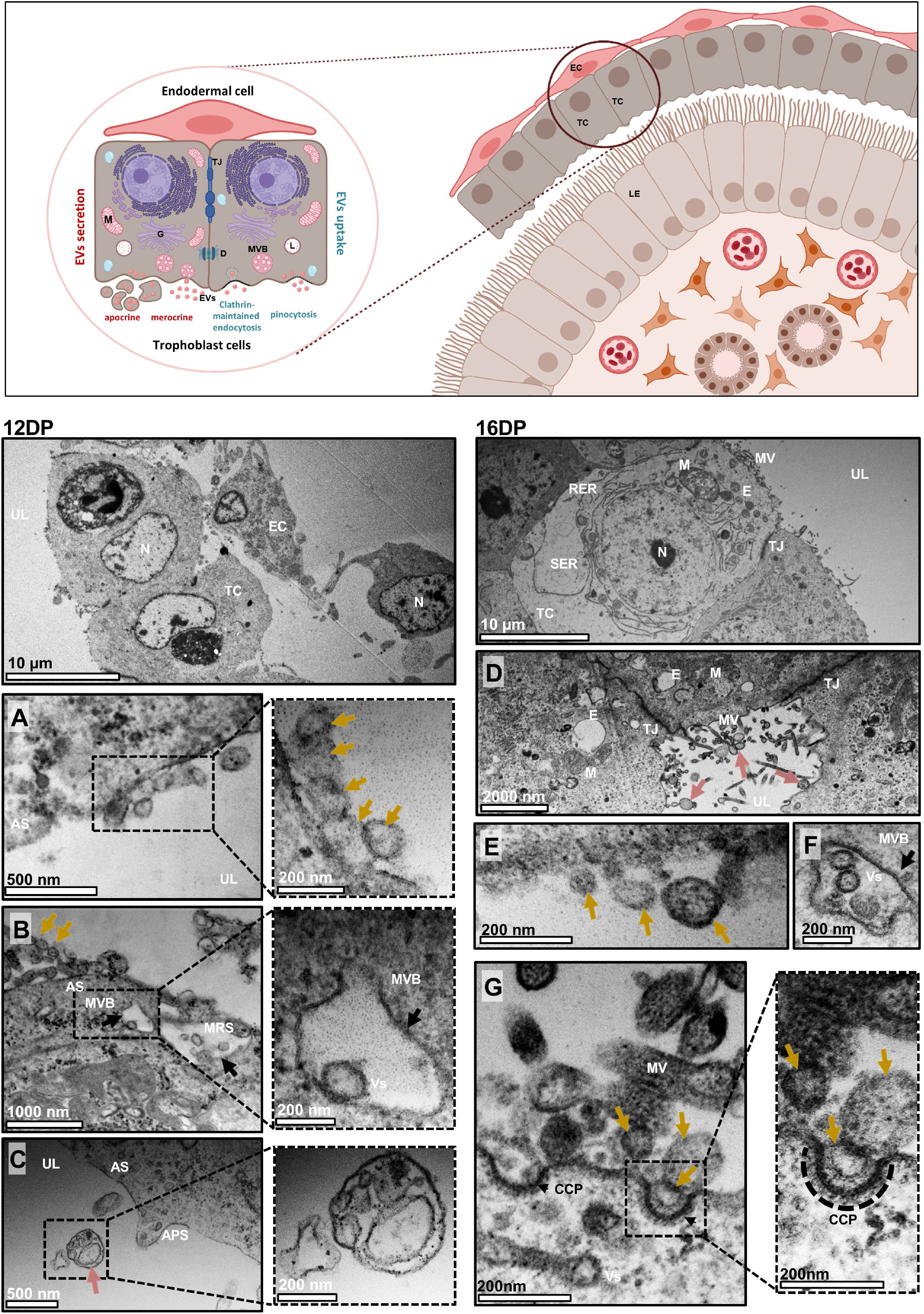
EVs are released by early embryos via apocrine and merocrine secretory mechanism. *Upper panel:* Schematic representation the embryo after apposition/attachment to the luminal epithelium. Trophoblast cells are facing the luminal epithelium to maintain the contact between the developing embryo and receptive endometrium. The basal cell surface of trophoblast cells stays in a close contact with elongated endodermal cells. Porcine trophoblast secretes EVs via apocrine (when apical part of the secretory cell containing vesicles pinches off and enter the into the uterine lumen) and merocrine (when vesicles are released from cell *via* exocytosis) mechanism. Trophoblast cells also receive molecules from the uterine environment. Clathrin-maintained endocytosis and pinocytosis are portals for uptake of molecules and particles, including EVs. *Left panel:* Cross section of the embryonic cells on day 12 of pregnancy. **(A)** Different in size and shape EVs (yellow arrows) covered by double membrane (higher magnification) were seen attached to the plasma membrane at the apical surface of trophoblast cell. **(B)** Merocrine type of secretion was evident at the apical surface of the trophoblast cells, exemplified by the fusion of numerous MVBs with plasma membrane (black arrows) and vesicles ready to be released into the extracellular space (higher magnification). **(C)** Apocrine type of secretion (pink arrows) was also observed, as the cytoplasmic fragments containing the numerous secretory vesicles. *Right panel:* Cross section of the trophectoderm on day 16 of pregnancy. **(D)** Micrograph showing close connections between trophoblast cells through tight junctions. Close to the apical surface of cells contain numerous secretory vesicles and endosomes. Apocrine type of secretion (pink arrows) was also observed. **(E)** EVs (yellow arrows) attached to the apical surface. **(F)** MVBs (black arrow) filled with vesicles were located under the cell membrane at the apical surface. **(G)** Clathrin-mediated endocytosis was observed. Clathrin-coated pits were formed at the apical surface of the trophoblast cell membrane. Double-membrane EVs were visible inside the clathrin-coated pits (higher magnification). AS – apical surface, APS – apocrine secretion, CCP – clathrin-coated pit, DP – day of pregnancy, EC – endodermal cell, E – endosomes, M – mitochondria, MRS – merrocrine secretion, MV – microvilli, MVB – multivesicula bodies, N –nucleus, RER – rough endoplasmic reticulum, SER – smooth endoplasmic reticulum, TC – trophoblast cell, TJ – tight junctions, UL – uterine lumen, Vs – vesicles.

On 12 DP and 16 DP, simple columnar epithelial cells were located on the basal lamina with condensed nuclei, and connected apically by desmosomes and tight junctions. On 12 DP, a heterogeneous population of free EVs was captured in the uteri lumen; docking on the microvilli was indicated (Fig. 1A). MVBs were observed at the apical site of the luminal epithelium (Fig. 1B). The same pattern of the secretion and localization of EVs was observed on 16 DP (Fig. 1D, E). Interestingly, for the first time, characteristic secretory areas (SA) between two luminal epithelial (LE) cells were observed at this time point. These areas contained inside a population of heterogeneously sized EVs (Fig. 1C).

Along the pregnancy progression, increasing secretory activity of glandular cells was indicated (12 vs. 16 DP; Fig. 2A, C). Furthermore, numerous EVs were found in the lumen of the glands and large MVBs were localized at the apical region of the glandular epithelial (GE) cells (Fig. 2B, C). The trophoblast cells changed in size and shape, from elongated, almost oval cells on 12 DP, to large cubic cells on 16 DP (Fig. 3; see upper low-magnification images for each DP). Additionally, we detected different types of EV secretory mechanisms in trophoblast and luminal epithelial cells. Merocrine secretion was commonly seen in both endometrium and trophoblast cells as a fusion between MVBs and the apical cell membrane (Fig. 1B, E; 2B; 3B, F), allowing the free release of single vesicles (Fig. 1A, D; 2B, C; 3A, E). Apocrine secretion, observed only in trophoblast cells, was exemplified by the shedding of whole pieces of the cytoplasm containing EVs (Fig. 3C). This type of secretion was evident during the intensive phase of embryo development on 12 DP. Several regions of trophectoderm were highly active in EV-mediated communication. Vesicles inside characteristic pits were located at the apical surface of the plasma membrane of trophoblast cells on 16 DP (Fig. 3G), suggesting clathrin-dependent or independent internalization.

To achieve a full view of the EV secretion patterns at the embryo–maternal interface during early pregnancy, we examined later stages of pregnancy (20 DP). We found that the EV-mediated secretory activity of luminal epithelial cells had decreased, as free vesicles docking to the smooth surface of the plasma membrane were rarely seen (Supplementary Fig. 3A, B). However, large MVB filled with numerous vesicles were still visible (Supplementary Fig. 3B). MVBs were also present in the apical regions of GE cells. Moreover, EVs released into the lumen of the gland were noticed (Supplementary Fig. 3C, D). Two types of EV secretory mechanisms were still noticed on 20 DP at the embryo–maternal interface, as in other examined days of pregnancy. Both free EVs (Supplementary Fig. 3E, G) released via the merocrine pathway and EVs released with cytoplasmic fragments (Supplementary Fig. 3F) via the apocrine mechanism were observed.

Overall, these data showed EV-mediated cell-to-cell communication between developing porcine embryos and endometrium (maternal interface) during early pregnancy.

### 3.2 Proteins involved in biogenesis and secretion of EVs show spatiotemporal pattern of distribution at the embryo–maternal interface

The contributions of EVs to cell-to-cell communication between the luminal/glandular epithelium and trophectoderm at early pregnancy stages suggest that the cellular localization of proteins involved in EV biogenesis and trafficking may change dynamically at the embryo–maternal interface. Thus, we selected two important members of the ESCRT complex network for further investigation—VPS37B, a subunit of ESCRT-I (MVB formation [16]), and Alix, an associated protein of the ESCRT pathway (the final step of EV biogenesis, crucial for MVB generation, [17]). Both were additionally colocalized with Rab27A (responsible for EV secretion and cell polarization, [20, 21]). Days 12 and 16 of pregnancy were selected as the respective time points of initial and more advanced implantation phases—before the final attachment to the luminal epithelium is completed, but after the embryonic signal has already been sent. Importantly, only untouched trophoblast–luminal epithelial interfaces were investigated to gain transparent insight into embryo–maternal interactions. Our results indicated spatiotemporal patterns of protein localization between 12 DP and 16 DP, showing close interaction between the luminal epithelium and developing embryo. On 12 DP, right after the embryonic signal occurred, Alix was highly stained in the trophoblast cells, as well as in the lumen/apical site of the epithelium of the uterine glands (Fig. 4A, B), while VPS37B was found only in the trophectoderm and scattered in the cytoplasm of the glandular epithelial cells (Fig. 5A, B). In contrast, during ongoing embryo–maternal interaction on 16 DP, Alix was readily expressed in luminal and glandular epithelial cells but exhibited low to no expression in the trophectoderm (Fig. 4C, D). On the other hand, the VPS37B signal in the trophectoderm was polarized apically toward the luminal epithelium and was missing in the uterine glands (Fig. 5C, D). Interestingly, Rab27A showed also different cellular distribution in endometrium and trophectoderm (Fig. 4, 5). On 12 DP, the expression of Rab27A in the luminal epithelial cells was lower than in the trophectoderm, but on 16 DP it was almost evenly detected on both sides of the dialogue. Rab27A seemed to govern vesicle-mediated transport at the trophoblast attachment sites rather than in the uterine glands, as its expression was rarely seen in the glandular epithelium. The results consistently showed that along with the progression of early pregnancy and occurrence of embryonic signaling, the biogenesis and trafficking of EVs change at the embryo– maternal interface.

**Figure 4.**
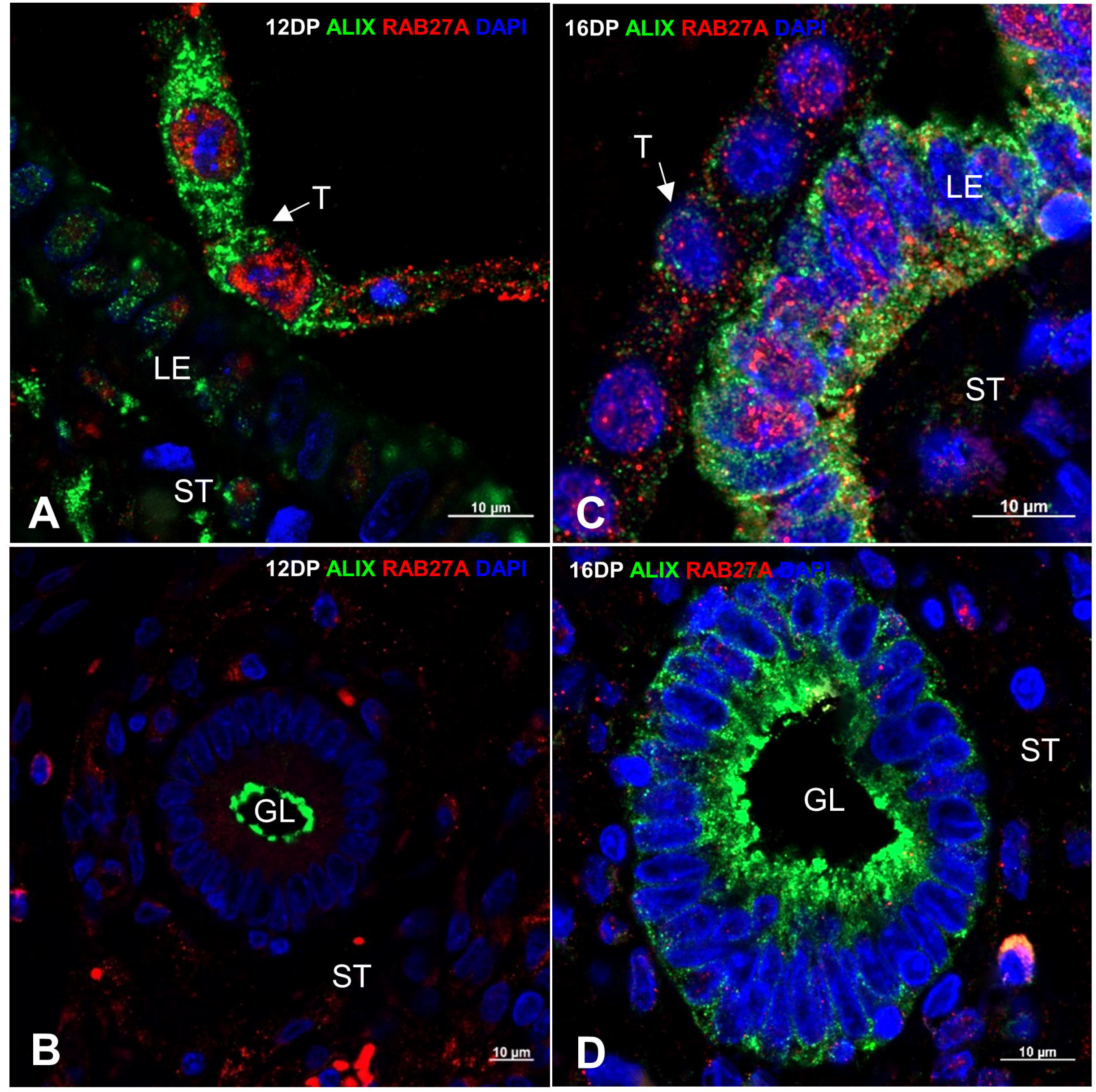
Spatiotemporal distribution of Alix and Rab27A at the embryo–maternal interface changes dramatically change along with the progression of pregnancy. Representative immunofluorescence images displaying the untouched trophoblast–luminal epithelial interfaces and colocalization of Alix (green) and Rab27A (red) proteins. Nucleus is depicted in blue (DAPI). Trophectoderm attached to uterine luminal epithelium on days 12 **(A)** and 16 **(C)** of pregnancy. Uterine glands surrounded by stromal cells on days 12 **(B)** and 16 **(D)** of pregnancy. GL – glands, LE – luminal epithelium, T – trophectoderm, ST – stromal cells.

**Figure 5.**
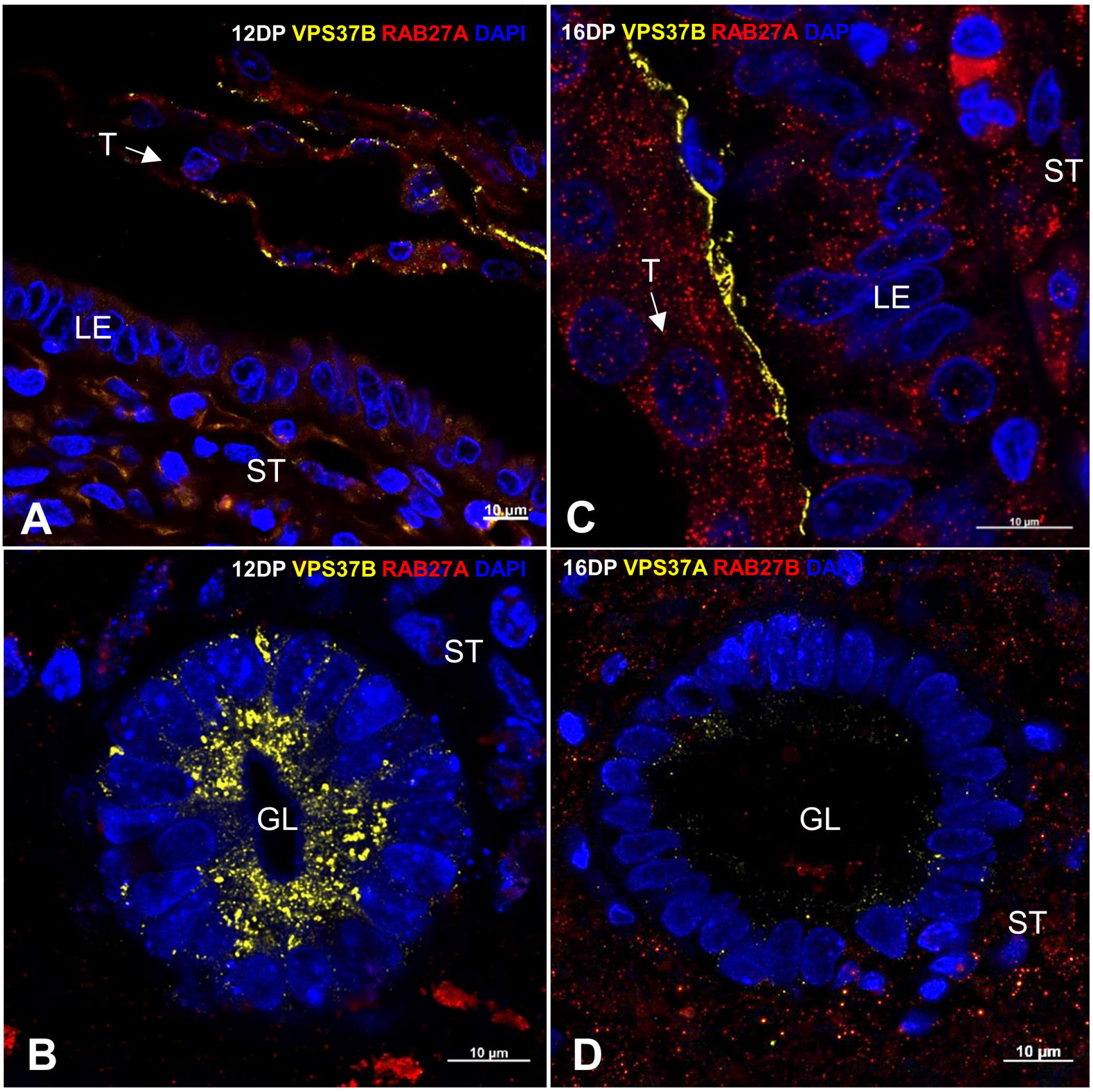
Spatiotemporal distribution of VPS37B and Rab27A at the embryo–maternal interface changes dramatically change along with the progression of pregnancy. Representative immunofluorescence images displaying the untouched trophoblast–luminal epithelial interfaces and colocalization of VPS37B (yellow) and Rab27A (red) proteins. Nucleus is depicted in blue (DAPI). Trophectoderm attached to uterine luminal epithelium on days 12 **(A)** and 16 **(C)** of pregnancy. Uterine glands surrounded by stromal cells on days 12 **(B)** and 16 **(D)** of pregnancy. GL – glands, LE – luminal epithelium, T – trophectoderm, ST – stromal cells.

### 3.3 Endometrial expression of genes coding EV biogenesis and secretion proteins change upon arrival of embryo into the uterus

To track changes in cell-to-cell communication in the endometrium evoked by the presence of embryos, we first analyzed the expression of genes involved in EV biogenesis and trafficking. *VPS28* and *VPS37B* were chosen in order to evaluate the expression of genes coding the ESCRT-I complex proteins responsible for the transport and sorting of cargo to EVs (Fig. 6A) [16]. *VPS28* expression in the endometrium was affected by both main factors (day and reproductive status). In the following days of the estrous cycle and pregnancy, the expression of *VPS28* decreased, while *VPS37B* showed almost constant expression. The second analyzed group of genes code proteins of ESCRT-II complex—*VPS22, VPS25*, and *VPS36*—which are all required for MVB formation (Fig. 6B) [32]. Expression of *VPS25* and *VPS36* was affected by the day, with increased expression on days 11–12 of the estrous cycle versus pregnancy. In the estrous cycle, *VPS22, VPS25*, and *VPS36* showed the highest expression on days 11–12, decreasing in the following days to reach the lowest levels on days 17–19. Stable levels of *VPS36* in the first days of pregnancy were observed, with a sharp drop after day 16. The third group of analyzed genes consisted of *Alix*, vesicle associated membrane protein 8 (*VAMP8*), and *VPS4A* (Fig. 6C). *Alix*, responsible for the last step of cytokinesis and intraluminal vesicle formation [33], was clearly affected by the day. *VAMP8*, involved in the secretion of EVs [34], and *VPS4A*,responsible for the redistribution of proteins belonging to ESCRT complex back to cytoplasm [12], showed stable expression during the estrous cycle and pregnancy (Fig. 6C). The last group included genes coding for RAB proteins involved in EV trafficking—*Rab11A, Rab11B, Rab7B*, and *Rab27A* (Fig. 6D) [20, 21]. The *Rab11A* expression was affected by reproductive status. Although its expression decreased gradually during the estrous cycle, it was maintained at a constant during pregnancy. On days 17–19, the expression of *Rab11A* was increased during pregnancy comparing to estrous cycle. Almost identical patterns of expression were observed for *Rab11b*. *Rab27A* expression was affected by day and the highest expression was observed on days 11–12 of the estrous cycle. On the other hand, *Rab7B* expression was affected by reproduction status, increasing in the consecutive days of pregnancy (Fig. 6D). These results support the conclusion that when an embryo arrives in the uterus, EV biogenesis and trafficking machinery is changed in the endometrium due to the presence of embryonic signals, as depicted in Figure 6E.

**Figure 6.**
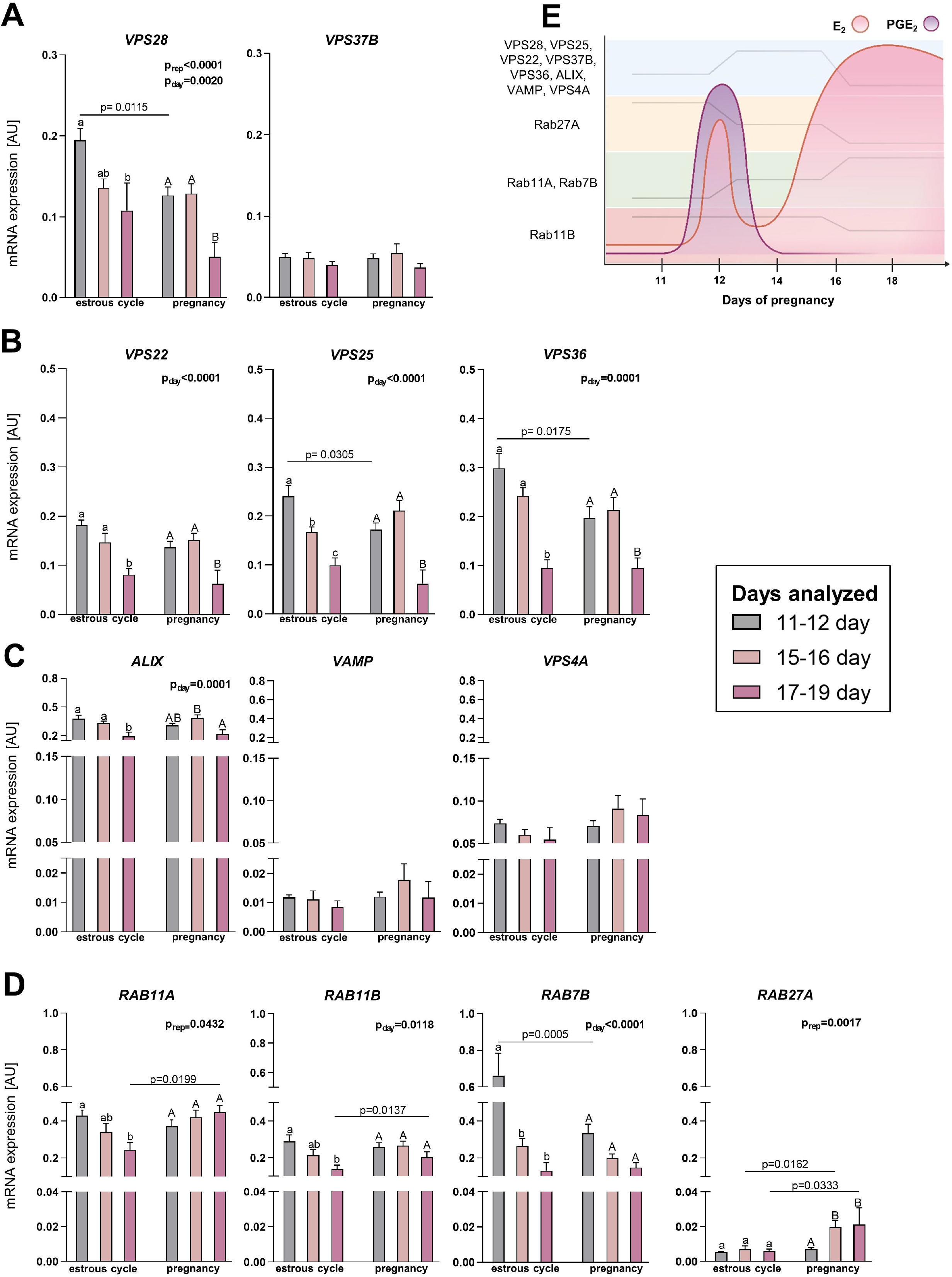
EV biogenesis and trafficking pathways in the endometrium are affected by the arrival of embryo in the uterus. mRNA expression levels of **(A)** ESCRT-I complex proteins: *VPS28*, and *VPS37b*, **(B)** ESCRT-II complex proteins: *VPS22, VPS25*, and *VPS36*, **(C)** ESCRT-associated proteins: *Alix, VAMP8*, and *VPS4a*, and **(D)** Ras-related proteins: *Rab11a, Rab11b, Rab7b*, and *Rab27a*. Gene expression was normalized to GAPDH (AU), identified as the best reference gene by NormFinder algorithm. Data were analyzed using two-way ANOVA with Sidak multiple comparison test and are shown as means + SEM (n = 5-13). Means with different superscripts differ significantly (small letters – the estrous cycle, capital letters – pregnancy; p < 0.05). Significant differences between the estrous cycle and pregnancy at the same day are indicated in the graph (p < 0.05). AU – arbitrary units. (E) Schematic diagram presenting combined gene expression and embryonic signals (E_2_ and PGE_2_) profiles during early pregnancy in pigs.

### 3.4 Binding motifs for estrogen and prostaglandin response elements are present in *VPS25* and *VPS36* promoter regions

As gene expression analysis suggested that the embryonic signaling may be involved in the modulation of EV biogenesis and trafficking, we tested whether E_2_ and PGE_2_ affected gene abundance in uterine luminal epithelial cells—the first cell layer receiving the signal from the embryo. We screened in silico promoter regions of genes affected by either day (VPS25, *VPS28, VPS36*, and *Rab27A*) or/and reproductive status (*VPS25* and *VPS36*) to identify ERE (GATCAANNNGACC) [35, 36] and PGREs (PGRE1: TGACCTTC; PGRE_2_: GTCCCTCA; PGRE3: AGTCCCTGC) [37]. Notably, ERE and PGREs were only identified for *VPS25* and *VPS36*,components of the ESCRT-II complex (Fig. 7A, B).

**Figure 7.**
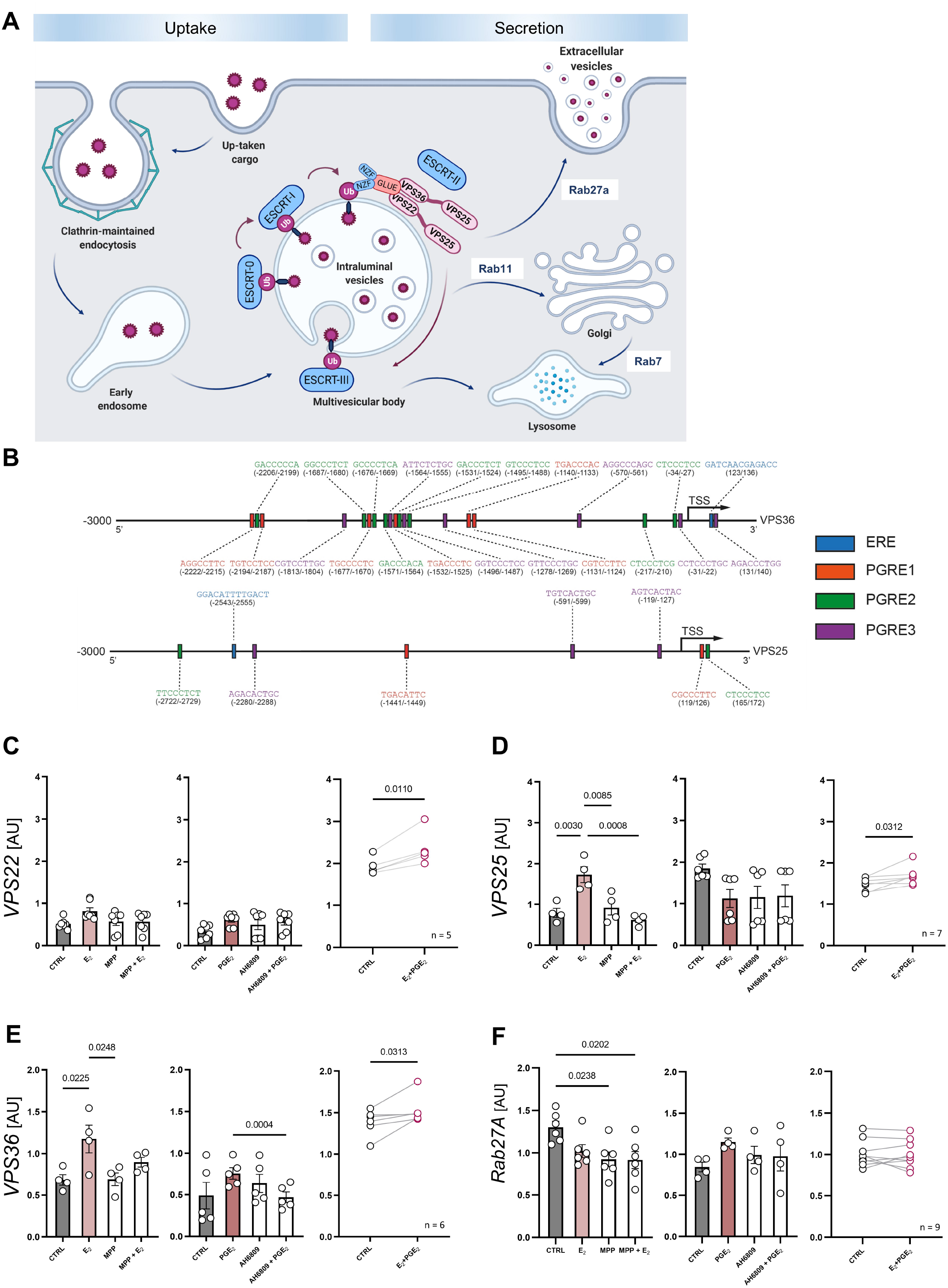
ESCRT-II complex in luminal epithelium is under the influence of embryonic signals. **(A)** Schematic representation of ESCRT complex and Ras-related proteins involved in EV biogenesis and trafficking. **(B)** Diagram showing possible positions of binding motifs for estrogen (ERE, blue box) and prostaglandin (PGRE1, red; PGRE_2_, green; PGRE3, purple boxes) response elements upstream/downstream from the VPS25 and VPS36 TSS. **(C–F)** mRNA expression of *VPS22, VPS25, VPS36*, and *Rab27a* in response to treatment with E_2_ (100 nM) or PGE_2_ (100 nM) in primary luminal epithelial cells. Cells were pretreated with specific inhibitors – MPP for ESR1 (1h, 1 μM) or AH6809 for PGE_2_ (10 minutes, 10 μM). Gene expression was normalized to HPRT1 (AU), identified as the best reference gene by NormFinder algorithm. Data for separate E_2_ and PGE_2_ treatments were analyzed using one-way ANOVA with Tukey’s multiple comparison test (*VPS25, VPS36, Rab27a*; D, E, F; p < 0.05) or Kruskal-Wallis and Dunn’s multiple comparisons test (*VPS22*; C), and are shown as means ± SEM (vs. control; n = 4 - 7). Data presenting simultaneous treatment with both embryonic signals were analyzed using paired T-test (C, D, and F; p < 0.05) or Wilcoxon matched-pairs signed rank test (E; p < 0.05) and are presented as individual plots (control-treatment). AU – arbitrary units, TSS – transcription start site.

### 3.5 Estradiol and PGE_2_ affect EV biogenesis and secretion pathways in luminal epithelial cells

Both VPS25 and VPS36 proteins are mainly required for MVBs formation, as members of ESCRT-II complex [32]. Thus, to see if ESCRT-II is indeed modulated by E_2_ and/or PGE_2_, we examined the expression of all genes coding proteins belonging to the ESCRT-II complex network—*VPS22*, *VPS25*,and *VPS36* (Fig. 7A). The impacts of the embryonic signal on the selected elements of EV biogenesis and transport pathways were tested in vitro using luminal epithelial cells collected on days 11–12 of the estrous cycle to assure the embryonic signal was in a naïve state in the primary cell culture. We found that *VPS25* and *VPS36* expression was stimulated by E_2_ (Fig. 7D, E), as suggested by the in silico analysis of ERE motifs in their promoter regions. Moreover, the pretreatment of luminal epithelial cells with the ESR1 antagonist (MPP) abolished the E_2_-mediated effect on gene expression (Fig. 7D). PGE_2_, the second embryonic signal tested, induced *VPS36* expression (Fig. 7E). The simultaneous exposition of luminal epithelial cells to both embryonic signals showed increased expression of both genes compared to the control (Fig. 7D, E). A different regulation pattern was observed for *VPS22*. Both embryonic signals added separately to the culture media showed no effect on *VPS22* gene expression. This observation was supported by the lack of ERE/PGREs motifs in *VPS22* promoter region. However, the simultaneous treatment (E_2_ and PGE_2_) slightly increased *VPS22* levels (Fig. 7C), suggesting that this component of the ESCRT-II complex is under different regulation requiring both embryonic signals. Notably, we were able to mimic in vitro the embryonic signal effects observed ex vivo, when decreased expression of *Rab27A* was indicated on days 11–12 of pregnancy compared to the estrous cycle (Fig. 6D). The expression of *Rab27A*, not involved in ESCRT-II complex formation and devoid of ERE/PGREs motifs in its promoter region, changed only for MMP and E_2_ combined with MPP (Fig. 7F). These results clearly indicate that both *VPS25* and *VPS36* genes, coding proteins of the ESCRT-II complex, are under the influence of embryonic signals during early pregnancy.

### 3.6 VPS36 and CD63 protein colocalization after embryonic signaling point to its triggering role in EVs trafficking in luminal epithelial cells

To further examine the impact of embryonic signaling on EV-mediated cell-to-cell communication, we focused on the protein localization in luminal epithelial cells exposed to embryonic signals. The origin of cell monolayers was proven by control staining with cytokeratin, as a marker of epithelial cells (Fig. 8A). VPS36 (marker of ESCRT-II complex present in MVBs, [32]) and CD63 (marker of EVs, [38]) were stained with specific antibodies after treatment with E_2_ and PGE_2_ added separately or simultaneously to luminal epithelial cell culture. We anticipated that depending on the dynamics of EV-mediated cell-to-cell communication, both proteins will be either colocalized in the same structure—MVBs filled with generated EVs—or not colocalized—for example, after the release of EVs from MVBs. Both proteins were detected in the cytoplasm and on the edges of luminal epithelial cells, indicating preserved cell-to-cell communication in vitro (Fig. 8B). The analysis of ROIs (marked in white; Fig. 8B) in the monolayers of the luminal epithelial cells (Fig. 8C), covering regions stained with both antibodies at the cell-to-cell contact areas, revealed specific patterns of VPS36- and CD63-stained puncta localization and their signal intensity. After E_2_ treatment, the fluorescence intensity of the VPS36 signal was significantly increased compared to the control group. Significantly, the intensity for the CD63 signal was more scattered after E_2_ delivery compared to the control, suggesting increased trafficking of EVs in the luminal epithelial cells (Fig. 8D). The colocalization coefficient analysis further pointed at dynamic changes in vesicular trafficking, showing differences between treatment groups in terms of colocalized pixels of both proteins CD63 and VPS36. In the controls, high colocalization coefficiency of CD63+ and VPS36+ was observed, meaning that almost all CD63+ puncta were present in the VPS36+MVBs. In the treatment groups, colocalization coefficiency of CD63+ and VPS36 decreased for all treatments (Fig. 8E). It suggests, accumulated number of CD63+ puncta not encapsulated in MVBs after treatment. Interestingly, the most striking observations were related to the VPS36+ and CD63+pixel colocalization correlation after treatment. In the control group, correlation between VPS36 and CD63 was high, followed by a characteristic linear signal distribution (Fig. 8F). In contrast, both were eliminated after E_2_ delivery to the cell culture. An almost zero distribution of signal reflects that VPS36 and CD63 were not related to one another after treatment. Similar, but not as pronounced, observations were captured for PGE_2_, as well as simultaneous treatment with E_2_ and PGE_2_. Observed changes in VPS36 and CD63 signal distribution between the control and treatment groups can be a significant premise of EV secretion after E_2_ and PGE_2_ delivery to the culture. These data suggest that embryonic signaling can affect vesicular trafficking in luminal epithelial cells.

**Figure 8.**
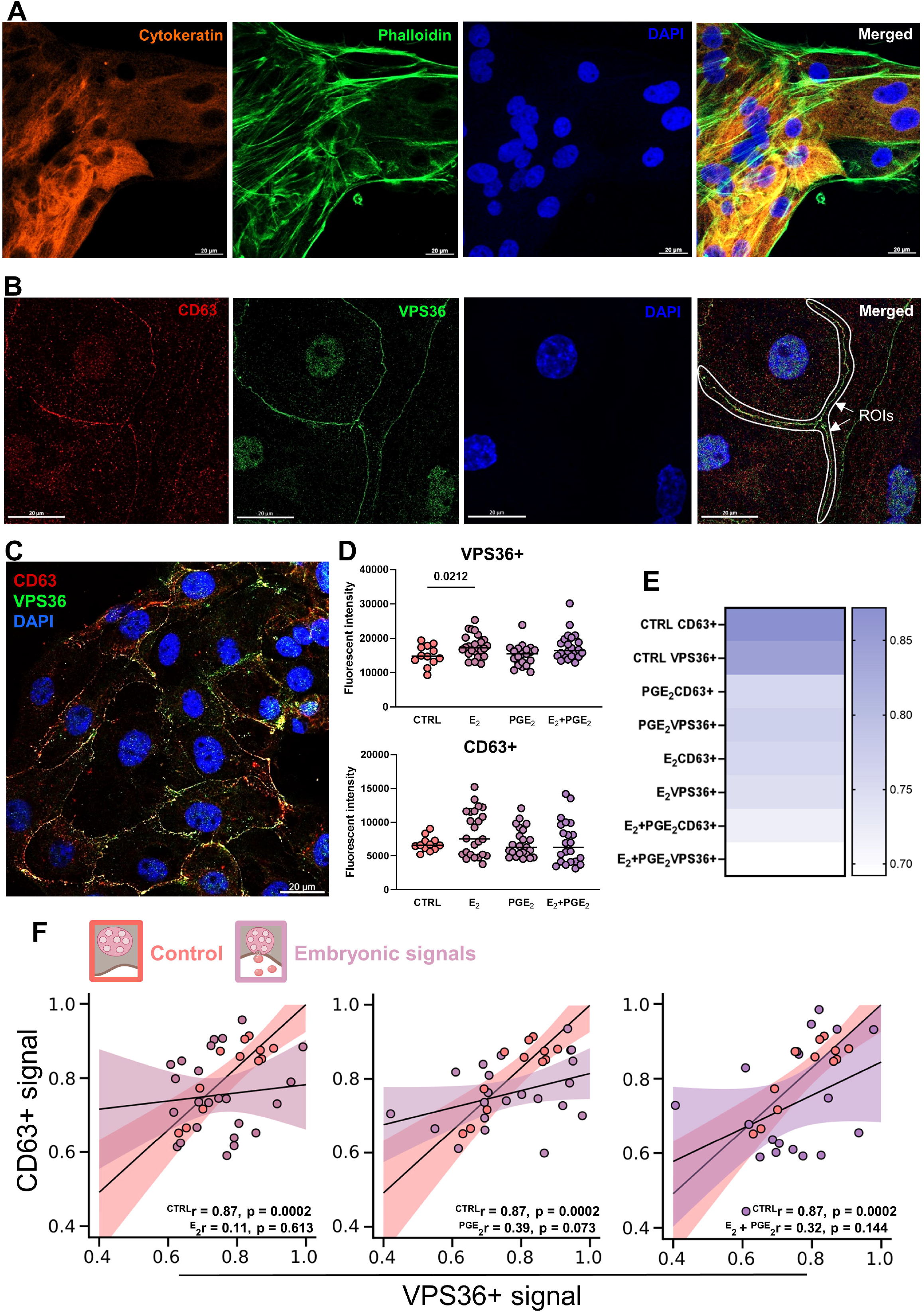
Embryonic signals affect vesicular trafficking in luminal epithelial cells. **(A)** Epithelial origin of primary cell culture was tested using cytokeratin marker. Actin filaments were marked with phalloidin. Nucleus was stained with DAPI. **(B)** Identification of regions of interest (ROIs) was performed based on the colocalization of CD63 (CY3, red) and VPS36 (Alexa 488, green) at the cell-to-cell contact areas. Nucleus was stained with DAPI. Merged images show an example of ROIs (white arrows). **(C)** A representative immunofluorescence image displaying luminal epithelial cell colony evaluated for intestines and colocalization. **(D)** Signal intensities for CD63 and VPS36 after treatment with embryonic signals - E_2_ or PGE_2_ (100 nM each; unpaired T-test, p < 0.05). **(E)** A heatmap of positively colocalized pixels of both proteins (Cy3 for CD63 and Alexa 488 for VPS36). Values are reported as pairs of pixels that range from 0 to 1 (no to full colocalization). **(F)** Scatter plots of CD63+ and VPS36+ pixels corresponding to the colocalization events as shown in B and C indicate correlation and linear relationship for control (r = 0.87, p = 0.0002) and lack of correlated and non-linear relationships for E_2_ (r = 0.11, p = 0.613), PGE_2_ (r = 0.39, p = 0.073) and a simultaneous treatment with E_2_ and PGE_2_ (r = 0.32, p = 0.114). Pearson correlation coefficient was performed (p < 0.05).

### 3.7. Embryonic signals have superior role over miRNAs in regulation of EV biogenesis and trafficking in luminal epithelial cells

In the final approach, relevance miRNA and hormone interplay in regulation of EVs biogenesis pathway was tested. First, we proved efficiency of miR-125b-5p delivery to cell culture by decreased expression of its target gene - *VPS36* (Supplementary Fig. 4A). Similar LE cells response was observed when miRNA was delivered after hormonal treatment (Fig. 9A). However, it was abolished when treatments were added in a reverse order (Fig. 9D). Based on described above method of ROIs analysis, we analyzed protein distribution after each sequence of treatments, focusing on cell borders and whole-cells. When miRNA was delivered after embryonic signals, VPS36+ and CD63+ pixels fluorescent intensities were different at cell borders in control and miRNA treatment (Fig. 9B), while whole cells followed the same pattern only in control (Fig. 9C). Also, low correlation between VPS36+ and CD36+ puncta was evident (Fig. 9B, C, lower panels). Embryonic signals delivered after miRNA led to increase of VPS36+ over CD63+ pixel fluorescence intensity at cell borders (Fig. 9E) and in whole cells (Fig. 9F), but more importantly remarkable VPS36+ signal increase was noticed at cell borders (Fig. 9E; Supplementary Fig. 4B). It was accompanied by a dramatic loss of correlation between VPS36+and CD36+ puncta (Fig. 9E, lower panel). The same patter, though less extreme, was observed for whole cells (Fig. 9F). This is a strong indication of specialized cell-to-cell communication at cell borders, governed by embryonic signals.

**Figure 9.**
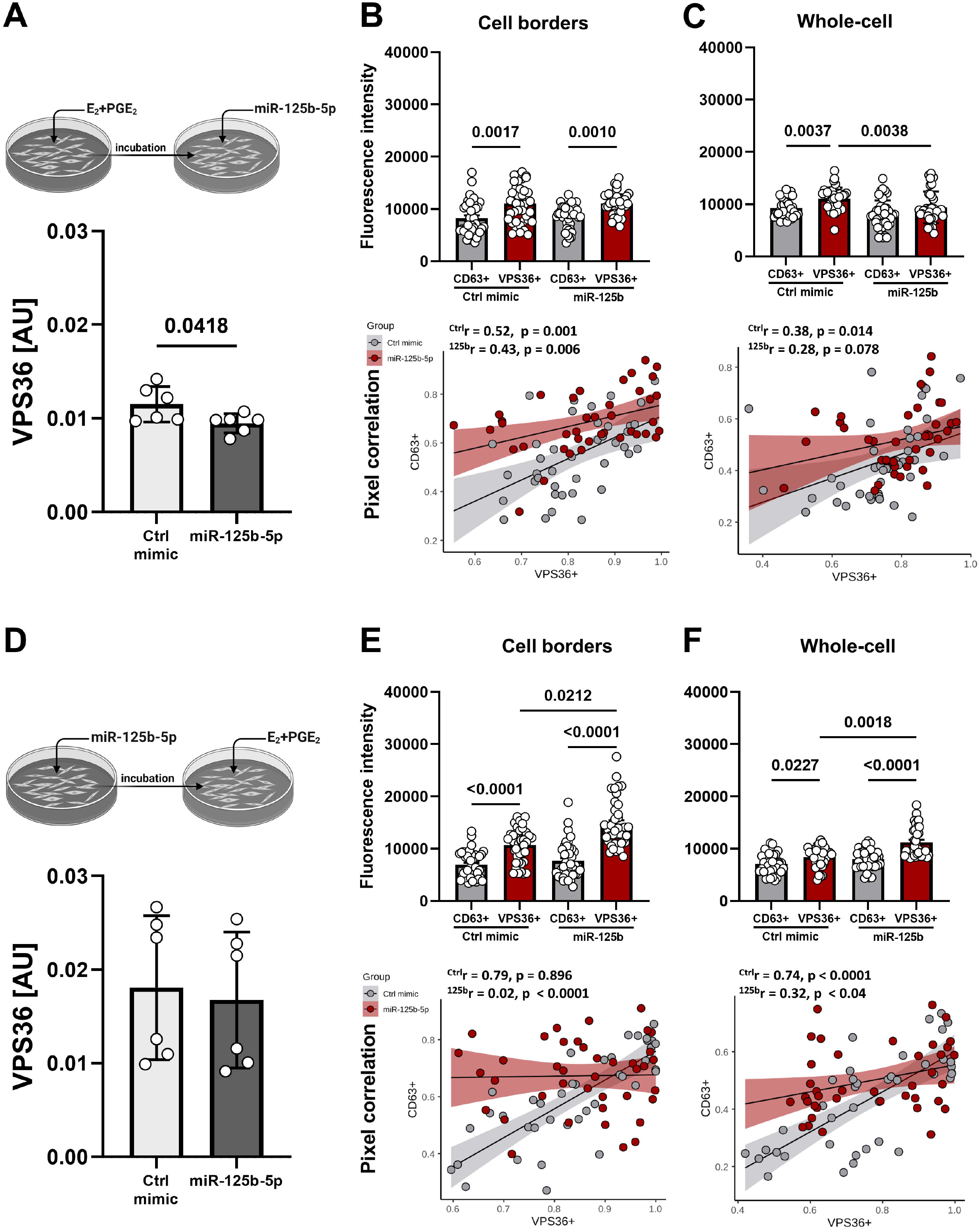
Embryonic signals override fast regulatory action of miRNA in governing EV biogenesis pathway in luminal epithelial cells. **(A)** Luminal epithelial cells were exposed to embryonic signals, E_2_ and PGE_2_ (24 h), followed by miR-125b-5p delivery (12 h) to test *VPS36* target gene expression (unpaired T-test). Gene expression was normalized to HPRT1 (AU), identified as the best reference gene by NormFinder algorithm. Signal intensities (upper panels) and pixel correlations (lower panels) for CD63 and VPS36 at cell borders **(B)** and in whole cells **(C)** for miRNA delivered after embryonic signals (one-way ANOVA followed by Kruskal-Wallis post hock test, Pearson correlation coefficient). **(D)** Luminal epithelial cells were transfected with miR-125b-5p (12 h), which was followed by exposition to embryonic signals, E_2_ and PGE_2_ (24 h), to test *VPS36* target gene expression (unpaired T-test). Gene expression was normalized to HPRT1 (AU), identified as the best reference gene by NormFinder algorithm. Signal intensities (upper panels) and pixel correlations (lower panels) for CD63 and VPS36 at cell borders **(E)** and in whole cells **(F)** for embryonic signals delivered after miRNA (one-way ANOVA followed by Kruskal-Wallis post hock test; Pearson correlation coefficient).

## 4 DISCUSSION

The role of EVs in cell-to-cell communication occurring during early pregnancy in mammals has been highlighted in many studies. Essential components carried by EVs, such as miRNAs and proteins, have attracted much attention in EV-mediated function studies; however, little attention has been paid to EV biogenesis and trafficking at the embryo–maternal interface. Here, we demonstrated diverse mechanisms of EV secretion and uptake in luminal epithelial and trophoblast cells during early pregnancy. During the embryo apposition and attachment phases, which are accompanied by release of pregnancy recognition signals, the spatiotemporal changes in the expression of molecules involved in EV biogenesis and trafficking were seen at the intimate contact areas between embryonic and maternal surfaces. Consistent with these findings, embryonic signaling for pregnancy recognition (predominantly E_2_ via ESR1) was found to affect the expression of genes coding proteins involved in EV biogenesis and change EV trafficking at the maternal site (endometrium, luminal epithelium), bringing about a novel regulatory mechanism of EV-mediated cell-to-cell communication at the initial stages of mammalian pregnancy.

Transmission electron microscopy confirmed that the luminal epithelium and trophoblast cells undergo morphological and functional changes during pregnancy, leading to the establishment of unique embryo–maternal communication. These observations are consistent with early reports that tight junctions between endometrial luminal epithelial cells become located in the basolateral region, the nuclei become larger, and the cytoplasm is less dense with glycogen droplets at the basal side [1, 13, 39–44]. Our results identified characteristic secretory areas filled with vesicles, pointing at an unseen earlier type of cell-to-cell communication occurring between two metabolically active luminal cells. This is strong evidence of horizontal communication occurring between luminal cells, which involves EVs in auto/paracrine intercellular transport. In addition, we confirmed the presence of either MVBs filled with EVs or free EVs on both sides of the initial embryo–maternal dialogue. Observed endocytosis was proposed as a mechanism through which EVs can actively interact with target cells and participate in intercellular communication [45, 46]. This type of gateway for the entry of extracellular substances into the cell was observed in trophoblast cells, confirming the involvement of EVs in communication between the embryo and the mother at early stages of pregnancy.

The ongoing vesicle-based communication at the initial stages of pregnancy was shown through the dynamic (in time and space) localization of proteins involved in EV synthesis and trafficking. Interactions between molecules within ESCRT complexes have been investigated first in yeast, and then in mammals, indicating that ESCRT complexes are broadly conserved among species [32, 47]. In addition, ESCRT complexes were found to regulate several aspects of cell migration and polarization [48–50], processes evident at the early stages of pregnancy in mammals. VPS37B, a component of the ESCRT-I complex, which mediates vesicular trafficking [51, 52], switched cellular localization from distracted on day 12 of pregnancy to highly polarized on day 16, but only in trophoblast cells at the implantation sites. Alix, as an accessory protein of ESCRT, tightly involved in EV cargo loading, endocytic membrane trafficking, and cytoskeletal remodeling [33, 49, 53], showed characteristic, distinct expression in the endometrium and trophoblast. Initially, Alix was predominantly present in the trophectoderm, but along with the initiation of a second wave of embryonic estrogen secretion (i.e., 15–30 days of pregnancy) [5], its expression switched to the luminal epithelium. Likewise, the expression patterns of Rab27A was depended on the cell type (luminal/glandular epithelium vs. trophectoderm) and the analyzed day of pregnancy. An intriguing finding was that on 12 DP, Rab27A was almost exclusively present in the trophoblast cells, shifting its expression to luminal epithelium on day 16. This suggests the involvement of Rab27A in gaining cell polarity during the development of close cell-to-cell contact between the trophoblast and uterine epithelium [20, 21]. Indeed, Rab27A was found to be involved in the regulation of epithelial cell polarity through EV-dependent mechanisms, leading to the intercellular reorganization of vesicles transported to the apical membrane for the extracellular delivery of proteins [54]. The polarization of the examined proteins in the trophectoderm toward the luminal epithelium, following the waves of embryonic signaling, hints at the relevance of EV-mediated interactions during the apposition and attachment of the trophectoderm to the uterine epithelium.

Estradiol plays a key role in maintaining pregnancy and reproductive functions [55]. Its cellular signaling occurs with subsequent binding to nuclear receptors ESR1 [56]. Aside from estrogens, PGE_2_ has been suggested as involved in the maternal recognition of pregnancy in pigs, as a luteoprotective/antiluteolytic factor acting predominantly through PTGER2 and PTGER4 [7]. The importance of embryonic signaling in the establishment and maintenance of pregnancy and our microscopic observations suggested the involvement of E_2_ and PGE_2_ in EV-mediated cross talk at the embryo–maternal interface. Indeed, four main profiles of endometrial genes involved in EV biogenesis and trafficking coincide with the occurrence of embryonic signals (Fig. 6E). Consistent with prior studies indicating that all ESCRT complex proteins provide a bridge for the formation of a link between the previous/next complex [57], we showed similar expression profiles for interacting molecules (i.e., *VPS22, VPS25, VPS28*, and *VPS36;* Fig. 6E). Interestingly, the same pattern of expression was noticed for *Rab27A*,closely involved in EV secretion (Fig. 6E) [49, 58].

The characterization of uterine responsiveness to embryonic signaling on day 12 of pregnancy, showed that the expression of ESR1 [58, 59], as well as PTGER2 and PTGER4 [7], increases in the luminal and glandular epithelium. In silico analysis revealed the direct binding of ESR and PGREs to the promoter regions of the *VPS25* and *VPS36* genes, implying that the ESCRT-II complex (a trilobal complex containing two copies of VPS25 and one of each VPS22 and VPS36 [46]) stays under the control of embryonic signals. In our in vitro model, the responsiveness of the naïve luminal epithelium to E_2_ was verified via ESR1 and PGE_2_ through PGTER2 (involved in a PGE_2_ positive feedback loop in the porcine endometrium [7]). Consistent with the bioinformatics predictions, E_2_ treatment induced the expression of *VPS25* and *VPS36* via ESR1 but not *VPS22*, the latter of which has a promoter devoid of ERE and PGRE binding motifs. Notably, the simultaneous administration of E_2_ and PGE_2_ treatment increased the expression of all ESCRT-II genes (*VPS22, VPS25*, and *VPS36*). These results support the concept that embryonic signals are able to affect expression of all components of the ESCRT-II complex. This interaction may be governed not only by a direct interaction between ESR1 and ERE but also by the alternative transcription pathways, such as an indirect binding with another transcription factor without direct DNA binding [56]. However, further work is required to demonstrate the existence of such interactions. Nevertheless, E_2_-mediated stimulation of *VPS25* expression, a dominant core protein of the ESCRT-II complex, aided by VPS36 induction can be sufficient to set the functioning of an entire complex [60]. E_2_ and PGE_2_ treatment did not change the expression of *Rab27A* in luminal epithelial cells, although *VPS25, VPS36*, and *Rab27A* followed the same pattern of expression in the endometrium during pregnancy (Fig. 6E). A possible explanation lies in the lack of ERE and PGRE binding motifs within the promoter region of the *Rab27A* gene, as well as the complex cellular composition of the endometrium not reflected in vitro. There is a chance that *Rab27A* gene expression is governed by different regulatory mechanisms in the particular cell type (e.g., luminal and glandular epithelium, stromal cells).

One of the most intriguing parts of this study was the discovery that E_2_ and PGE_2_ may direct vesicular trafficking in the uterine luminal epithelium, the first line of cells receiving a signal from embryos present in the uterine cavity. Based on the E_2_-mediated induction of the ESCRT-II complex required for MVB formation [46] in the luminal epithelium, we decided to track MVBs and EVs after the delivery of embryonic signals to the naïve luminal epithelium. We found that the overlapping occurrence of VPS36 (a marker of ESCRT-II in MVBs, [52, 60]) and CD63 (a marker of EVs, [38]) in the cell-to-cell contact areas was diffused after E_2_ and PGE_2_ treatment. This suggested that embryonic signaling elicits EVs secretion from the MVBs/luminal epithelium, although this warrants further investigation. It is particularly important to understand how embryonic signals modulate EV-mediated endometrial remodeling at early stages of pregnancy to allow for embryo apposition and adhesion.

Our previous studies pointed at important regulatory function of miRNAs during early pregnancy [12, 28, 61]. Apparently, miR-125b-5p, an important gene expression regulator at embryo-maternal interface, occurred also as a negative regulator of VPS36. Thus, we decided to use miR-125b-5p in testing miRNA-embryonic signal interplay in regulation of EVs biogenesis and trafficking. Strikingly fast action of miRNA on VPS36 expression and VPS36+ and CD63+ signal distribution in luminal epithelial cells, especially at cell borders, was abolished when embryonic signaling occurred. This results strengthen our conclusion on pivotal role of embryonic signals in governing EV-mediated cell-to-cell communication. However, important role of miRNAs present in the uterine environment should be also noticed, as fast-acting regulatory molecules of EVs biogenesis and release during endometrial response to embryonic signals.

In summary, using both ex vivo and in vitro models, we have provided novel insights into the role of embryo–maternal signaling in governing EV biogenesis and trafficking pathways at the embryo–maternal interface during the early stages of mammalian pregnancy. In a combined microscopic approach, we detected and characterized EV-based features of early cell-to-cell communication occurring between endometrial epithelial cells and the trophectoderm. We have shown that the timely polarization of either trophoblast or epithelial cells to generate and release EVs coincides with the apposition and attachment phases of embryo implantation and the release of signals for the maternal recognition of pregnancy. Moreover, our results provide the first direct evidence that embryonic signals modulate the biogenesis and trafficking of vesicles in the luminal epithelium, the first line of cells receiving signaling from the developing embryo. Here, we propose a new regulatory mechanism of EV biogenesis and trafficking pathways at the embryo–maternal interface, leading to the reprogramming of the endometrium during early pregnancy, as a path toward communication coordinated in time and space between the developing embryo and receptive uterus.

## ACKNOWLEDGEMENTS

The authors are grateful to M. Romaniewicz, M. Sikora, B. Szerejko, K. Gromadzka-Hliwa, M. Blitek, J. Klos, Dr D. Panas and Dr K. Witek from the Institute of Animal Reproduction and Food Research, Polish Academy of Science for either excellent technical assistance laboratory or support in statistical data analysis; M. Angenitzki from The Hebrew University of Jerusalem for invaluable help with electron microscopy imaging. The study was supported by grant funded by KNOW (Leading National Research Centre) Scientific Consortium “Healthy Animal - Safe Food” (decision of Ministry of Science and Higher Education No. 05-1/KNOW2/2015; grant number - KNOW/2016/IRZiBZ/PRO1/01/5 to MMK), National Science Center in Poland (2018/29/N/NZ9/02331 to MMG) and Israel Science Foundation (ISF-2041/17 to YH). Scientific staff exchange was supported by Polish-Israel Joint Research Projects (2017-2019) under the agreement on scientific cooperation between the Polish Academy of Science and the Israel Academy of Science and Humanities. Graphics were created with BioRender.com.

## AUTHOR CONTRIBUTIONS

MMG designed and performed experiments, collected, analyzed, and interpreted data; prepared graphics, drafted the manuscript and participated in the preparation of its final version. MMG and KM contributed to the bioinformatics analyses. YH was responsible and supervised the transmission electron microscopy imaging, and participated in the preparation of the final version of manuscript. MMK conceived and supervised the study, designed experiments, analyzed, and interpreted data and was responsible for the final version of the manuscript.

## DATA AVAILABILITY STATEMENT

All data generated or analyzed during this study are included in this published article or in the supplementary material.

## ETHICAL STATEMENT

All procedures involving animals were conducted in accordance with the national guidelines for agricultural animal care in compliance with EU Directive 2010/63/UE.

## Supporting Information

Supplemental material is available for this paper.

