## Supporting Information for "Embryonic signals mediate extracellular vesicle biogenesis and trafficking at the embryo–maternal interface"

Figures

Supplementary Figure 1. Control staining performed without primary antibodies.

Supplementary Figure 2. mRNA expression in luminal epithelial cells.

Supplementary Figure 3. Ultrastructural changes in luminal/glandular epithelium and trophectoderm on day 20 of pregnancy.

Supplementary Figure 4. miRNA effect on target gene and protein abundance.

Tables

Supplementary Table 1. List of primary and secondary antibodies used in immunofluorescent staining of endometrial/trophoblast tissue and luminal epithelial cells.

Supplementary Table 2. List of mimics used in miRNA and hormone treatment experiment.

Supplementary Table 3. List of genes and TaqMan gene expression assays used in real-time PCR.

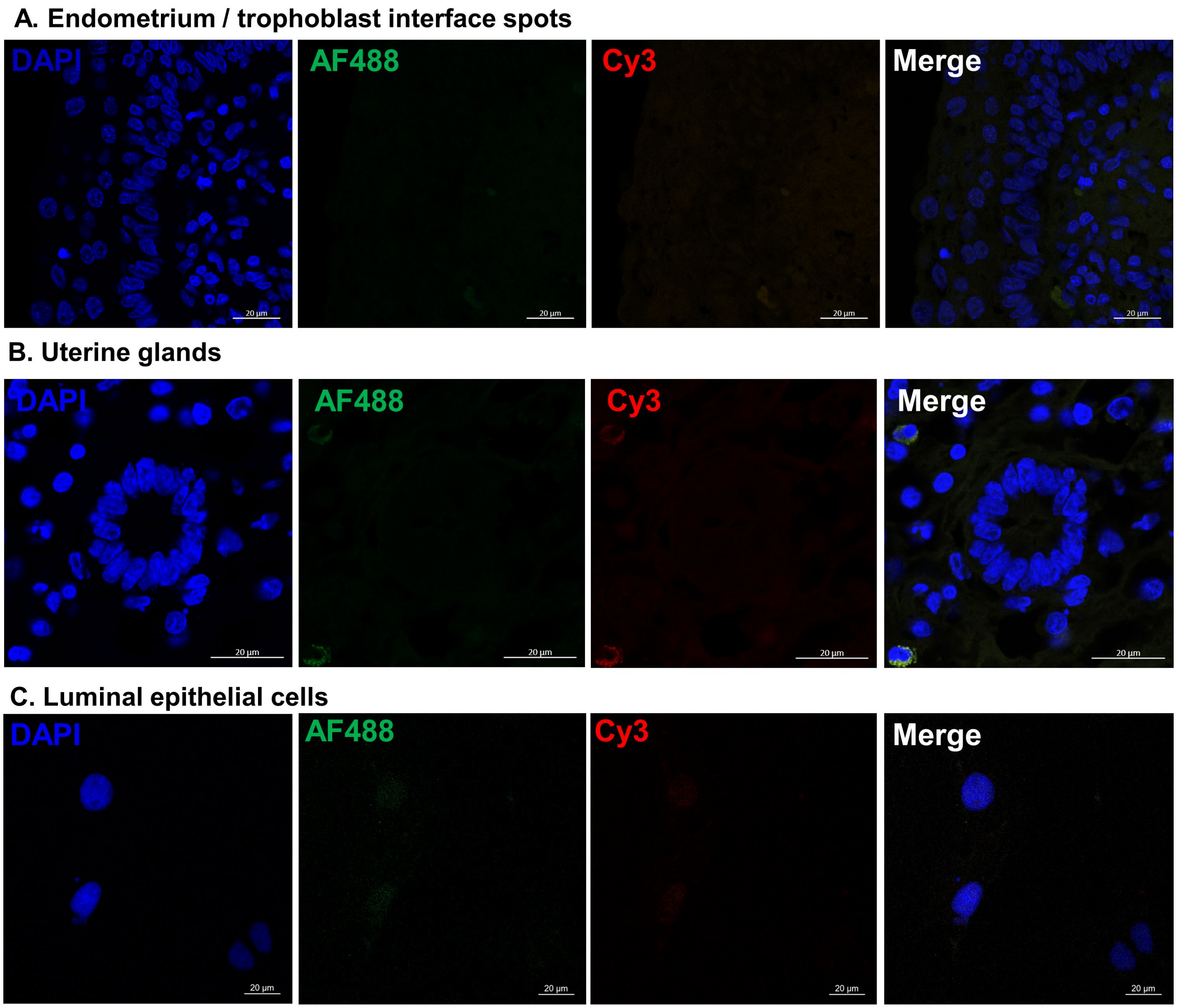


**Supplementary Figure 1.** Control staining performed without primary antibodies. **(A)** Negative control for untouched trophoblast-luminal epithelial interfaces. **(B)** Negative control for uterine glands. **(C)** Negative control for luminal epithelial cells. Merged pictures with red (Cy3-conjugated antibodies), green (Alexa Fluor 48-conjugated antibodies) and blue (DAPI) staining.

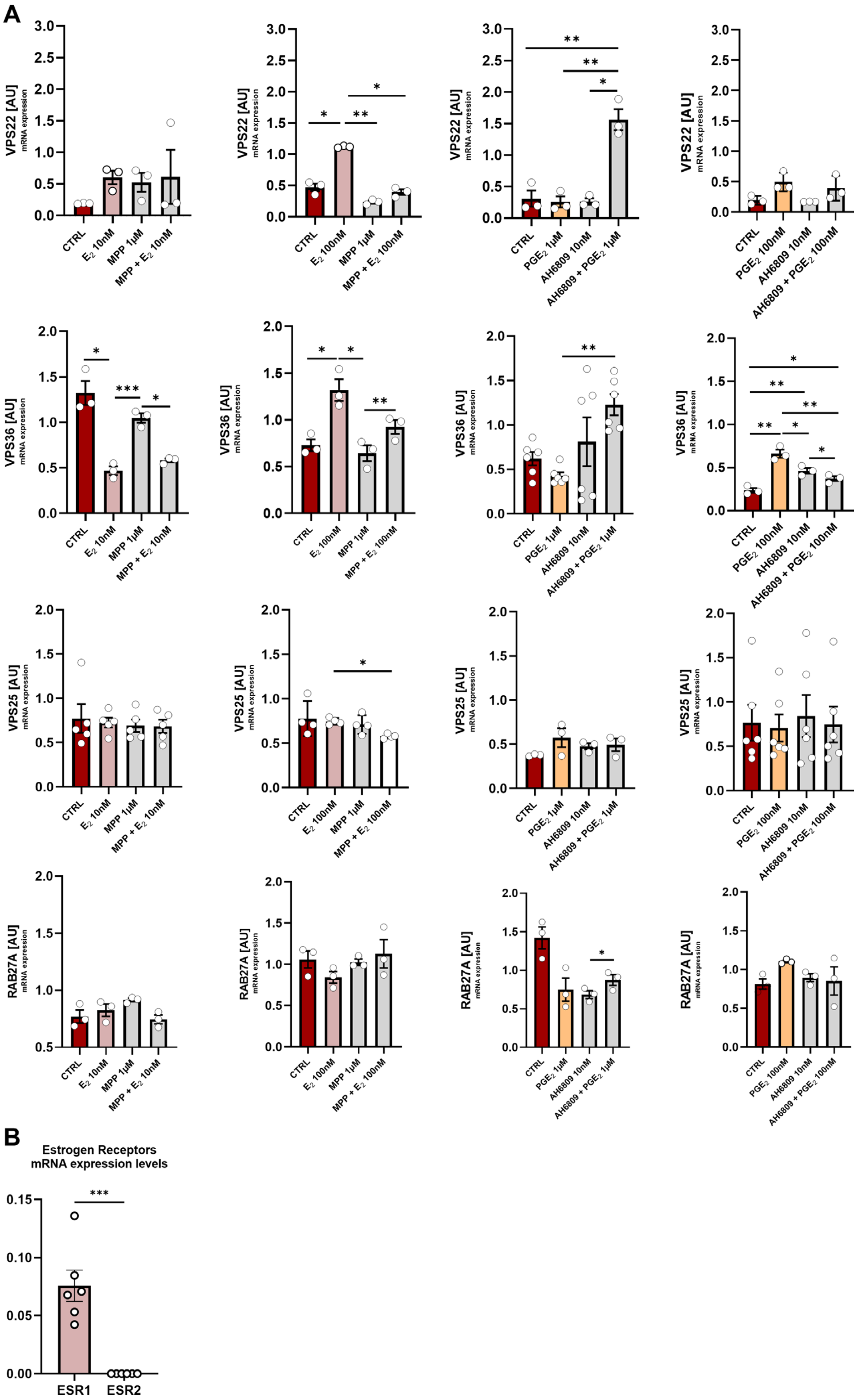


**Supplementary Figure 2.** mRNA expression in luminal epithelial cells. **(A)** VPS22, VPS25, VPS36, and Rab27A gene expression in response to treatment with E2 or PGE2 of primary luminal epithelial cells. Different doses of E2 (10 nM or 100 nM, pink boxplots) and PGE2 (1 µM or 100 nM, yellow boxplots) were used. Cells were pretreated with specific inhibitors – MPP for ESR1 (1h, 1 µM) or AH6809 for PGE2 (10 minutes, 10 µM). Data were analyzed using one-way ANOVA with Tukey's multiple comparison test (vs. control; * $p < 0.05$, ** $p < 0.01$, *** $p < 0.001$) and are shown as means ± SEM (n = 3 - 6). **(B)** ESR1 and ESR2 gene expression in response to treatment with E2 (100 nM). Data were analyzed using unpaired T- test (*** $p < 0.001$) and are shown as means ± SEM (n = 6). mRNA expression was normalized to HPRT1 (AU), identified as the best reference gene by NormFinder algorithm.

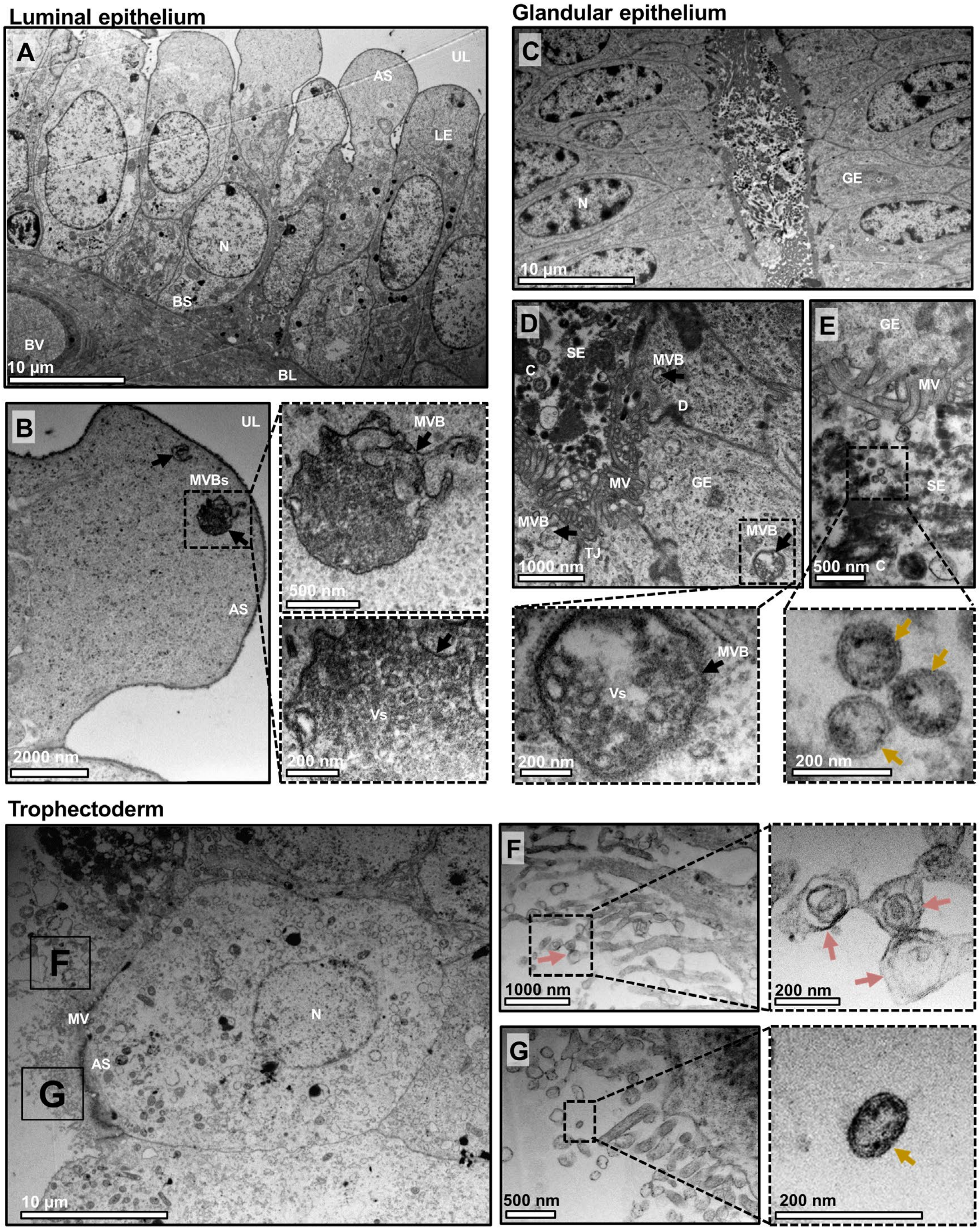


**Supplementary Figure 3.** Ultrastructural changes in luminal/glandular epithelium and trophectoderm on day 20 of pregnancy. **(A)** Cross section of luminal epithelium showing tall, columnar epithelial cells with large nucleus and any organelles at the apical site of the cell. **(B)** Large MVB about to fuse with plasma membrane were noticed at the apical site. MVBs contained heterogeneous population of double-membrane vesicles ready to be released into the extracellular space (higher magnification). **(C)** Cross section of wide open uterine gland. Large population of glandular cells with cilia, microvilli were observed. Glands were filled with secretome, containing heterogeneous population of EVs (yellow arrows), cached between the microvilli and cilia. The apical site of the glandular epithelial cells showed MVBs (black arrow) filled with double-membrane vesicles located under the plasma membrane (higher magnification). Lower panel: Cross sections of the trophectoderm. **(F)** Apical surface is covered with numerous microvilli. Apocrine type of secretion (pink arrows) was also observed, as the cytoplasmic fragments containing the numerous secretory vesicles (higher magnification). (G). Secreted, free double-membrane vesicles were found between the microvilli. AS – apical site, BL – basal lamina, BS – basal site, BV – blood vessels, C – cilia, D – desmosomes, GE – glandular epithelium, LE – luminal epithelial cell, MV – microvilli, MVBs – multivesicular bodies, N – nucleus, SE – secretome, TJ – tight junctions, UL – uterine lumen, Vs – vesicles

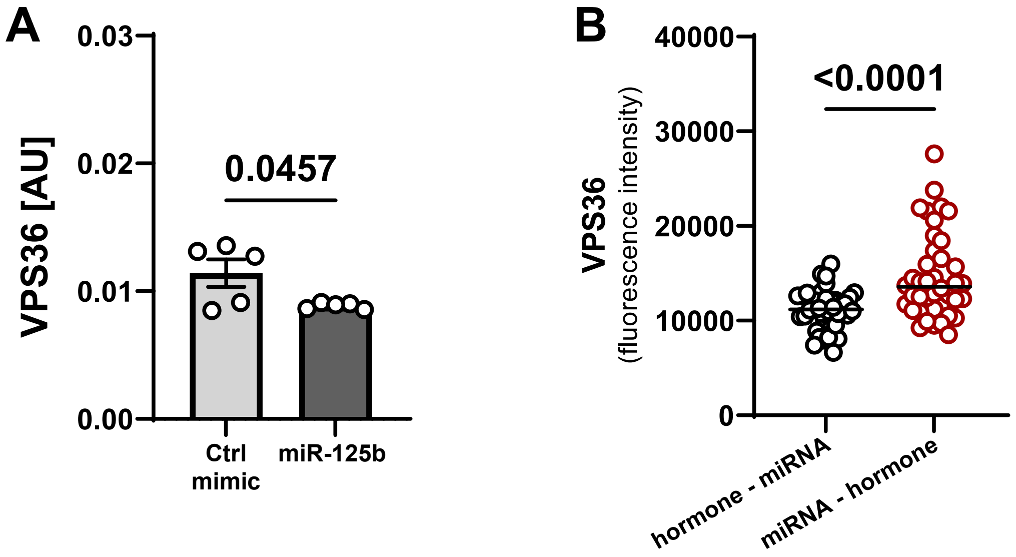


Supplementary Figure 4. miRNA effect on target gene and protein abundance. **(A)** miR-125b-5p affect the expression of target gene - VPS36 (p = 0.0457, unpaired T- test) in luminal epithelial cells. Data are presented as boxplots with means ± SEM. mRNA expression was normalized to HPRT1 (AU), identified as the best reference gene by NormFinder algorithm. **(B)** Fluorescence intensity of VPS36 signal in cell borders after miR-125b-5p delivery to luminal epithelial cells. Comparison between treatments revealed significant increase of VPS36+ puncta when embryonic signals were delivered after miRNA (p < 0.0001, unpaired T-test).

.

**Supplementary Table 1.** List of primary and secondary antibodies used in immunofluorescent staining of endometrial/trophoblast tissue and luminal epithelial cells.

| | Primary antibodies | Host | Description | Dilution | Catalog number | Company |
|---|---|---|---|---|---|---|
| TISSUE ENDOMETRIUM/ TROFOBLAST | Anti - VPS37B | Rabbit | Polyclonal | 1:50 | ab122450 | Abcam |
| | Anti - Rab27A | Mouse | Monoclonal | 1:50 | ab55667 | Abcam |
| | Anti - Alix | Rabbit | Polyclonal | 1:50 | PA5-52873 | Invitrogen |
| LUMINAL EPITHELIAL CELLS | Anti - VPS36 | Rabbit | Polyclonal | 1:50 | PA5-58905 | Invitrogen |
| | Anti - CD63 | Mouse | Monoclonal | 1:50 | ab193349 | Abcam |
| | Anti - Cytokeratin | Mouse | Monoclonal | 1:50 | C-9687 | Sigma |

| Secondary antibodies | Host | Conjugated dye | Dilution | Catalog number | Company |
|---|---|---|---|---|---|
| Anti-Mouse IgG | Donkey | Cy3 | 1 : 3000 | 715-165-150 | Jackson Immuno-Research |
| Anti - Rabbit IgG | Donkey | Cy3 | 1 : 3000 | 715-165-152 | Jackson Immuno-Research |
| Anti - Rabbit IgG | Goat | Alexa Fluor 488 | 1 : 3000 | A-21206 | Invitrogen |

**Supplementary Table 2.** List of mimics used in miRNA and hormone treatment experiment.

| miRNA mimic | Assay name | Assay ID | Mature miRNA Sequence | Catalog number |
|---|---|---|---|---|
| **miR-125b-5p** | hsa-miR-125b-5p | MC10148 | UCCCUGAGACCCUAACUUGUGA | 4464067 |
| **control mimic** | *mir* Vana™ miRNA Mimic, Negative Control #1 | | | 4464059 |

Supplementary Table 3. List of genes and TaqMan gene expression assays used in real-time PCR.

| Gene symbol | Gene name | Group/family/complex name | Assay ID |
|---|---|---|---|
| *Rab11A* | Rab11a | RAS Oncogene Family | Ss03379468_u1 |
| *Rab11B* | Rab11B | RAS Oncogene Family | Ss04322113_m1 |
| *Rab27A* | Rab27A | RAS Oncogene Family | Ss03392039_m1 |
| *Rab7B* | Rab7B | RAS Oncogene Family | Bt03251067_m1 |
| *Rab8B* | Rab8B | RAS Oncogene Family | Bt03243268_m1 |
| *SNF8 (VPS22)* | Vacuolar-Sorting Protein SNF8 homolog | ESCRT-II Complex Subunit | Bt03245176_m1 |
| *VPS25* | Vacuolar-Sorting Protein 25 homolog | ESCRT-II Complex Subunit | Hs01003777_gH |
| *VPS28* | Vacuolar-Sorting Protein 28 Homolog | ESCRT-I Complex Subunit | Hs01598026_m1 |
| *VPS36* | Vacuolar-Sorting Protein 36 homolog | ESCRT-II Complex Subunit | Bt03262017_m1 |
| *VPS37B* | Vacuolar-Sorting Protein 37 homolog | ESCRT-I Complex Subunit | Ss04324043_m1 |
| *VPS4A* | Vacuolar-Sorting Protein 4 homolog A | ESCRT-associated protein | Ss04324243_m1 |
| *PDCD6IP (Alix)* | Programmed Cell Death 6 Interacting Protein | ESCRT-associated protein | Hs00994345_m1 |
| *VAMP8* | Vesicle Associated Mmbrane Protein 8 | ESCRT-associated protein | Bt03225817_m1 |
| *ESR1* | Estrogen Receptor 1 | Nuclear Receptor Subfamily 3 Group A | Ss03383398_u1 |
| *ESR2* | Estrogen Receptor 2 | Nuclear Receptor Subfamily 3 Group A | Ss03391479_m1 |
| *ACTB* | Beta-actin | Reference gene | Ss03376081_u1 |
| *GAPDH* | Glyceraldehyde 3-phosphate Dehydrogenase | Reference gene | Ss03375435_u1 |
| *HPRT1* | Hypoxanthine–guanine Phosphoribosyltransferase 1 | Reference gene | Ss03388274_m1 |
